# Interorganelle competition for linoleic acid underlies steatotic liver pathology

**DOI:** 10.64898/2026.01.11.698890

**Authors:** Chuanhai Zhang, Dengbao Yang, Hiroyuki Suzuki, Gonçalo Dias do Vale, Jingxuan Chen, Amogh Vaidya, Kamran Melikov, Alexandra Swisher, Jingjing Wang, Mengchen Ye, Jin Zhou, Qiyu Zeng, Meijuan Bai, Mei-Jung Lin, Jeon Lee, Hao Zhu, Daniel J. Siegwart, Yujin Hoshida, Jeffrey G. McDonald, Xing Zeng

**Author notes:** These authors contributed equally (Chuanhai Zhang, Dengbao Yang).

## Abstract

Lipid droplets (LDs) are traditionally viewed as protective organelles that sequester potentially cytotoxic lipids. However, whether and how LD biogenesis in pathological contexts actively rewires interorganelle lipid homeostasis to drive organelle dysfunction and disease progression remains unexplored. Here, we identify the adipocyte-enriched protein calsyntenin 3β (CLSTN3B) as a critical promoter of metabolic dysfunction-associated steatotic liver disease (MASLD). CLSTN3B, an ER-LD contact protein previously shown to support LD maturation in adipocytes, is robustly induced in mouse hepatocytes by peroxisome proliferator-activated receptor γ (PPARγ) in response to dietary caloric overload. CLSTN3B drives LD biogenesis and neutral lipid storage by stabilizing hemifusion-like ER-LD membrane bridges via its arginine-rich segment. These bridges preferentially recruit cone-shaped linoleoylated phosphatidic acid (PA), diverting linoleic acid (LA) into triacylglycerides (TAGs) rather than mitochondrial cardiolipin (CL), leading to disrupted cristae architecture, deficient ETC supercomplex assembly, elevated electron leak, and oxidative stress. Hepatocyte-specific CLSTN3B deletion impairs LD formation, reduces TAG accumulation, enhances fatty acid oxidation, restores CL maturation, and mitigates oxidative stress, collectively attenuating MASLD progression. Consistently, hepatic CLSTN3B expression correlates with fibrosis severity and progression in human MASLD. These findings position LDs as active regulators of interorganelle lipid partitioning and establish CLSTN3B as a key determinant of mitochondrial vulnerability, providing a general framework for how dysregulated organelle interfaces shape disease.

## Introduction

Lipid droplets (LDs) are ubiquitous and dynamic organelles that serve as intracellular depots for neutral lipids, primarily triacylglycerides (TAGs) and sterol esters. LDs are bounded by a phospholipid monolayer and originate from the cytoplasmic face of the endoplasmic reticulum (ER)(*1*). During subsequent expansion, LDs continue to receive phospholipids and proteins from the ER(*2*). Additional cytoplasmic proteins are later recruited to the LD surface, often through amphipathic motifs that recognize packing defects in the phospholipid monolayer, thereby promoting LD maturation(*3, 4*). While LDs are the defining feature of adipocytes and underlie their unique capacity to store excess energy, extensive LD biogenesis in non-adipocytes has been documented in a variety of pathological contexts, such as in central nervous system (CNS) cells in neurodegenerative diseases(*5, 6*), in cardiomyocytes in cardiomyopathy(*7–9*) and hepatocytes in metabolic dysfunction-associated steatotic liver disease (MASLD). It is generally accepted that LD-mediated lipid compartmentalization mitigates lipotoxicity by sequestering bioactive fatty acids and derivatives in inert forms(*10–12*). Nevertheless, beyond these protective effects, whether and how LD biogenesis may actively contribute to cellular dysfunction and disease progression in those pathological contexts remains a critical unresolved question.

MASLD, formerly known as non-alcoholic fatty liver disease (NAFLD), is now recognized as the most prevalent chronic liver disorder worldwide, affecting more than 25% of the global population(*13, 14*), and may represent a uniquely tractable system to probe the pathogenic consequence of abnormal LD biogenesis. MASLD encompasses a spectrum of liver pathology that begins with the accumulation of neutral lipids, mostly TAGs, in hepatocytes (steatosis) and can progress to inflammatory steatohepatitis, fibrosis, cirrhosis, and ultimately hepatocellular carcinoma(*15, 16*). While the initial stage of steatosis is considered relatively benign, the transition to more advanced stages of liver damage is clinically significant and remains incompletely understood(*17*). Accumulating evidence suggests that excessive LD biogenesis may actively drive this transition. For example, the I148M variant in patatin-like phospholipase domain-containing protein 3 (PNPLA3), one of the strongest genetic risk factors for MASLD, impairs TAG hydrolysis by sequestering α/β hydrolase domain-containing protein 5 (ABHD5, also known as CGI-58), thereby inhibiting adipose triglyceride lipase (ATGL)-mediated lipolysis(*18–22*). Conversely, loss-of-function mutations in the cell death-inducing DFFA-like effector protein B (CIDEB) protein, a protein localizing to ER and LD, confer protection against liver disease(*23, 24*). Genetic or pharmacological perturbation of CIDEB improves MASLD condition in mice by markedly reducing LD volume and enhancing fatty acid oxidation (FAO)(*25–28*). Similarly, silencing perilipin 2 (PLIN2), a major LD-coating protein, or diacylglycerol O-acyltransferase 2 (DGAT2), the enzyme responsible for TAG synthesis and driving LD biogenesis, both mitigate MASLD phenotypes(*29–33*). Together, these findings support the notion that excessive LD biogenesis exacerbates MASLD progression.

While the genetic evidence implicating LD biogenesis in MASLD progression is compelling, the molecular mechanism linking excessive LD biogenesis to hepatocyte damage is multifaceted, with mitochondrial dysfunction and oxidative stress emerging as central players. Structural and functional mitochondrial abnormalities, such as disrupted cristae organization, reduced ATP synthesis, and elevated reactive oxygen species (ROS) production, have been reported in MASLD patients(*34*). Oxidative stress-induced release of hepatocyte-derived damage-associated molecular patterns (DAMPs) is thought to serve as a key trigger that activates hepatic stellate cells (HSCs) into extracellular matrix-producing myofibroblasts and drives fibrosis progression(*17*). Notably, a recent study demonstrated that cardiolipin (CL), an inner mitochondrial membrane (IMM)-specific phospholipid essential for maintaining cristae architecture and supporting electron transport chain (ETC) function, is markedly reduced in both human MASLD samples and mouse models(*35*). Hepatocyte-specific CL deficiency in mice recapitulates core pathological features of human MASLD, suggesting a causal link between CL loss and mitochondrial dysfunction(*35*). These findings raise an intriguing possibility that excessive LD biogenesis in hepatocytes perturbs lipid metabolic homeostasis and secondarily compromises mitochondrial CL biosynthesis, providing a potential mechanistic framework for how steatosis drives mitochondrial dysfunction and liver injury. Nonetheless, the molecular events connecting LD biogenesis and mitochondrial CL perturbation remain to be elucidated.

Recent studies in adipocytes identified calsyntenin 3β (CLSTN3B) as a novel protein localizing to ER/LD contact sites via an N-terminal LD-targeting domain and a C-terminal ER transmembrane region(*36, 37*). Through an arginine (R)-rich motif that interacts with the cone-shaped phospholipid phosphatidic acid (PA), CLSTN3B promotes hemifusion-like membrane bridging between the ER and LD, facilitating ER-to-LD lipid trafficking required for LD expansion(*36*). In adipocytes, loss of CLSTN3B reduces LD phospholipid coverage, increases basal lipolysis, and limits lipid storage capacity(*36*). When placed on a chow diet, CLSTN3B expression is largely restricted to mouse adipocytes(*38*), consistent with its role in supporting the maturation of uniquely large LDs characteristic of adipocytes. However, hepatocytes in MASLD similarly acquire large LDs, raising the possibility that CLSTN3B may play a previously unrecognized role in pathogenic hepatic LD expansion and disease progression.

Here, we demonstrate that dietary caloric overload induces hepatic CLSTN3B expression via peroxisome proliferator-activated receptor γ (PPARγ) activation. CLSTN3B-driven LD biogenesis is facilitated by enrichment of cone-shaped linoleoylated PA at ER/LD membrane bridges, where they are converted into TAGs and become sequestered in LDs. This process occurs at the expense of mitochondrial linoleoylated phospholipids required for CL remodeling, thereby impairing mitochondrial function and driving MASLD pathology. Genetic or pharmacological depletion of CLSTN3B restores mitochondrial function, increases fatty acid oxidation, and mitigates steatosis and liver injury. Using hepatocytes as a model system, our findings identify CLSTN3B as a molecular link between pathological LD biogenesis and mitochondrial lipid insufficiency, providing a mechanistic framework for how aberrant LD expansion and interorganelle lipid competition can compromise organelle function and drive cellular pathology.

## Results

### Dietary caloric overload induces CLSTN3B expression in mouse hepatocytes

Given the close relationship between excessive LD biogenesis and MASLD pathogenesis, we compared the proteome of LD fractions isolated from mice fed a chow diet versus a Western diet (WD, 21.1% fat, 41% sucrose, 1.25% cholesterol by weight) to identify factors potentially involved in interorganelle communication. To this end, we filtered our dataset using previously published high-confidence LD proteomes and curated lists of mitochondria-LD and ER-LD contact-associated proteins(*36, 39, 40*). This analysis identified CLSTN3B, a protein previously considered adipocyte-specific and known to localize to ER-LD contact sites, as the most strongly induced protein under the WD condition (Fig. 1A).

**Figure 1.**
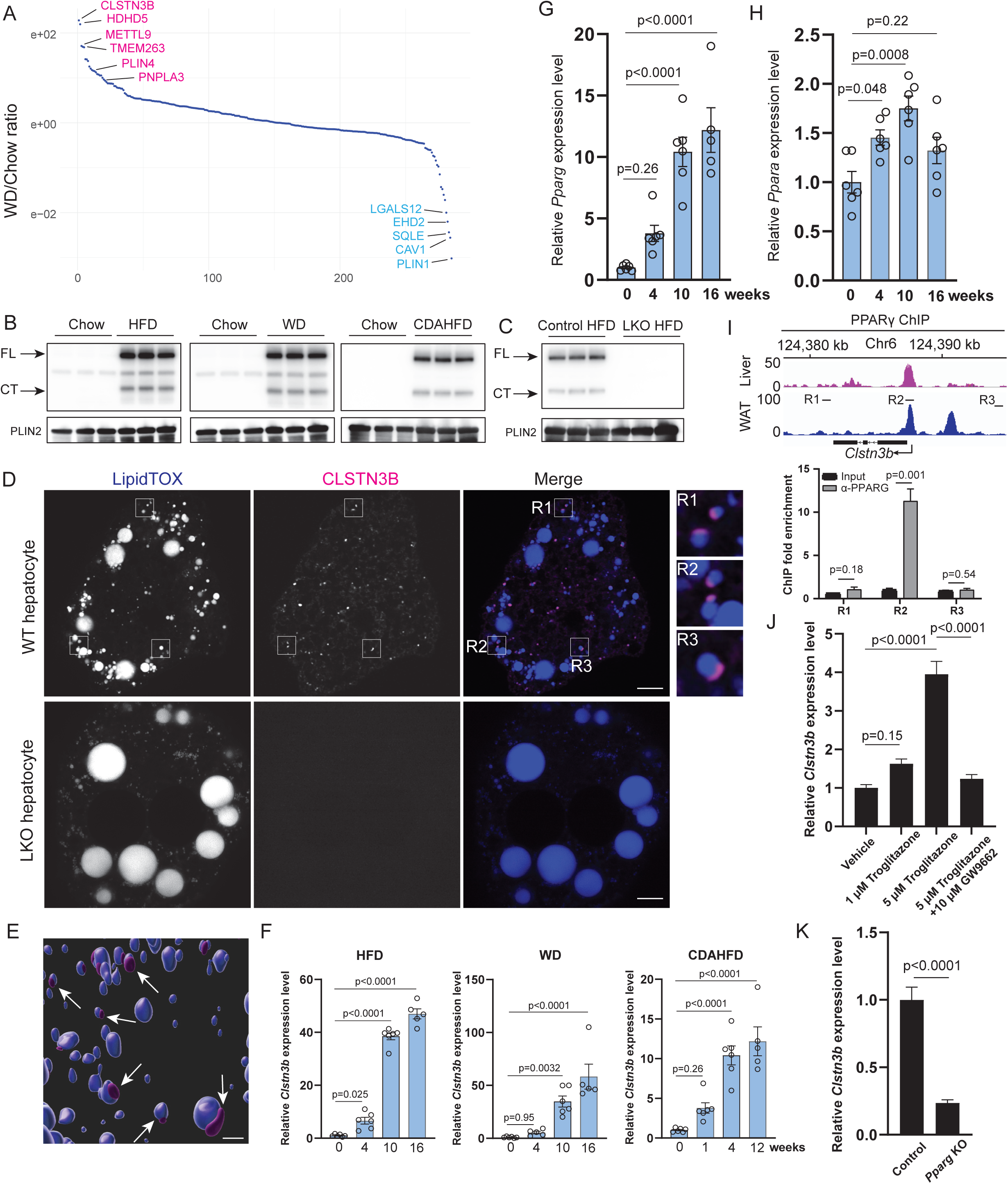
Dietary caloric overload induces CLSTN3B expression in mouse hepatocytes. **A**, Proteomics analysis of liver LD fractions isolated from mice on chow or WD for 10 weeks (n = 3 mice). **B**, Immunoblot analysis of CLSTN3B expression on liver LDs isolated from mice on different diets. **C**, Immunoblot analysis of CLSTN3B expression on liver LDs isolated from control or LKO mice on HFD. **D**-**E**, Fluorescence microscopic images (**D**) and 3D reconstruction (**E**) showing endogenous CLSTN3B localization in hepatocytes. **F**, qPCR analysis of liver *Clstn3b* expression from mice on different diets (n = 6 mice). **G**-**H**, qPCR of liver *Pparg* (**G**) and *Ppara* (**H**) expression from mice on HFD (n = 6 mice). **I**, Liver PPARγ ChIP-seq and ChIP-qPCR analyses identifying PPARγ binding sites upstream of the *Clstn3b* locus (n = 3 biological replicates). WAT PPARγ ChIP-seq at the same locus is shown as a comparison. **J**, Effect of PPARγ ligands on *Clstn3b* expression in hepatocytes (n = 4 biological replicates). **K**, qPCR analysis of liver *Clstn3b* expression from control or *Pparg* KO mice on HFD (n = 6 mice). Scale bar: 5 μm in **D** and 1 μm in **E**. Data are mean ± s.e.m. Statistical significance was calculated by one-way ANOVA with Tukey’s post hoc test (**F**, **G**, **H**, and **J**), or unpaired Student’s two-sided t-test (**I** and **K**).

Immunoblotting of hepatocyte LD fractions isolated from mice on fat-enriched diets, including a high-fat diet (HFD, 60% fat), a WD, or a choline-deficient high-fat diet (CDAHFD, 60% fat) for 12 weeks, all confirmed robust CLSTN3B induction (Fig. 1B), consistent with the proteomic findings. As previously reported in adipocytes(*36*), full-length CLSTN3B, and to a lesser extent, a C-terminal fragment, were detected in LDs from mice fed with the high-fat diets, but not from chow-fed controls (Fig. 1B). Both species were absent in hepatocyte LDs isolated from the liver-specific *Clstn3b* knockout (KO) mice (Fig. 1C) (LKO: *albumin-cre*; *Clstn3b^fl/fl^*) on HFD. Fluorescence microscopy revealed that endogenous CLSTN3B localizes to the LD surface in hepatocytes isolated from wild-type (WT) control and is absent in the LKO hepatocytes (Fig. 1D-E).

The diet-induced CLSTN3B expression happened at the transcription level because hepatic *Clstn3b* mRNA level increased markedly over time in response to the dietary interventions (a 40-50 fold induction after 16 weeks of HFD or WD, and a 12-fold induction at 12 weeks of CDAHFD, Fig. 1F). Previous studies have shown that high-fat feeding induces expression of PPARγ, a nuclear receptor critical for adipogenesis, in hepatocytes, where it transcriptionally activates adipocyte-selective genes(*41–43*). We found that *Pparg* expression in the liver is induced in mice on HFD following a similar time course to *Clstn3b* (Fig. 1G). A recent report showed that hepatic *Clstn3b* expression is induced ∼3-fold upon fasting in a peroxisome proliferator-activated receptor alpha (PPARα)-dependent manner(*44*). Compared to *Pparg*, *Ppara* transcript was only modestly induced by short-term HFD feeding, and the difference from baseline became insignificant by 16 weeks (Fig. 1H), suggesting that PPARγ may be the dominant regulator of HFD-induced expression of *Clstn3b* in hepatocytes. To test this, we performed PPARγ ChIP-qPCR from hepatocytes and identified a PPARγ binding site upstream of the *Clstn3b* transcription start site positioned similarly in white adipocytes (Fig. 1I). Treatment of hepatocytes with troglitazone, a PPARγ full agonist, significantly induced *Clstn3b* mRNA level in hepatocytes, an effect abolished by co-treatment with GW9662, a covalent non-agonistic PPARγ ligand (Fig. 1J). Finally, we induced liver-specific ablation of *Pparg* in mice on HFD by delivering AAV8-sgRNA targeting *Pparg* into Cas9-expressing mice after 2 weeks of doxycycline administration in drinking water. This resulted in a 72% reduction of hepatic PPARγ protein levels and an ∼80% reduction of *Clstn3b* transcript levels (Fig. 1K, Fig. S1A-D). Taken together, these findings establish PPARγ as a dominant transcriptional driver of *Clstn3b* expression in hepatocytes upon dietary caloric excess.

### Hepatocyte CLSTN3B deficiency impairs LD biogenesis and reduces lipid deposition

We previously showed that CLSTN3B employs an R-rich segment to induce the formation of hemifusion-like membrane bridges between the ER cytosolic leaflet and LD monolayer to facilitate LD biogenesis, thereby enhancing LD biogenesis and lipid storage in adipocytes(*36*). Given that substantial LD biogenesis happens in steatotic hepatocytes, we reasoned that the diet-induced CLSTN3B expression in hepatocytes may represent an adaptation (or maladaptation) to the increased demand for lipid storage. To test this idea, we analyzed male LKO mice and WT control littermates after 10 weeks of HFD feeding, starting at 6-8 weeks of age. Compared to WT controls, the LKO mice displayed significantly reduced liver mass, liver-to-body weight ratio, and liver TAG content (Fig. 2A-C). Gross liver appearance and histological analysis further confirmed markedly reduced lipid accumulation in LKO livers (Fig. 2D). In contrast to numerous LDs accumulated in WT hepatocytes, CLSTN3B-deficient hepatocytes tend to harbor a smaller number of substantially larger LDs (10-20 μm in diameter, see the inset of Fig. 2D and 2E-F). Phospholipid quantitation and proteomics analysis revealed decreased LD surface phospholipid coverage (Fig. 2G-H), increased abundance of class II perilipins, and decreased abundance of multiple class I proteins that access LDs via ER-LD membrane bridges (Fig. 2I, Supplementary Table 1). These findings suggest that LDs formed in the absence of CLSTN3B have high surface tension and more extensive coalescence, consistent with the role of CLSTN3B-mediated ER-to-LD phospholipid diffusion. Similar phenotypes were recapitulated in male LKO mice fed a WD (Fig. S2A-F) and, to a lesser extent, in female LKO mice on HFD, where the overall steatotic response was milder (Fig. S2G-K), consistent with the known sex differences in diet-induced hepatic lipid accumulation(*45*). AAV-induced deletion of *Clstn3b* yielded exactly the same phenotype as the constitutive genetic KO model (Fig. S2L-O), ruling out developmental compensation in the constitutive LKO model.

**Figure 2.**
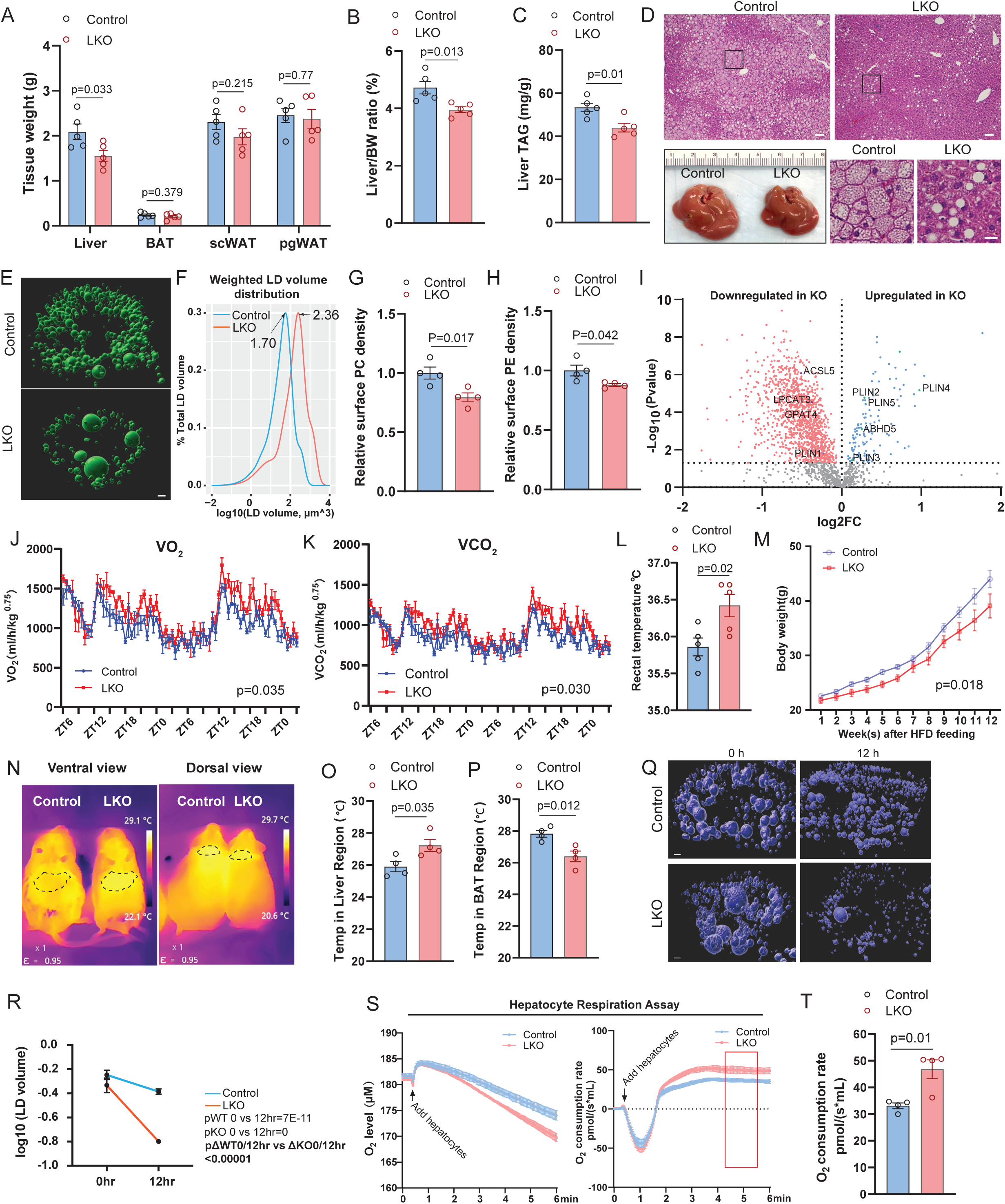
Liver specific deficiency of CLSTN3B alleviates HFD-induced hepatic steatosis. **A**-**D**, Tissue weights (**A**), liver/body weight ratio (**B**), liver TAG content (**C**), representive gross liver images and histology (**D**) from control and LKO mice on HFD for 10 weeks (n = 5 mice). **E**, 3D reconstruction of LD fluorescence images from control and LKO hepatocytes. **F**, weighted LD volume distribution curve for control and LKO hepatocytes. **G**-**H**, Quantitation of surface PC (**G**) and PE (**H**) densities on LDs isolated from control and LKO mice liver (n = 4 mice). **I**, Proteomics analysis of hepatocyte LDs isolated from control and LKO mice (n = 4 mice). **J**-**K**, Indirect calorimetry analysis of control and LKO mice on HFD for 10 weeks (n = 5 mice), showing O_2_ consumption rate (**J**) and CO_2_ production rate (**K**) normalized to body weight raised to the 0.75 power over a 45-hour period. **L**, Rectal temperature of control and LKO mice at ZT18 (n = 5 mice). **M**, Body weight growth curves of control and LKO cohorts maintained on HFD (n = 5 mice). **N**-**P**, Representative infrared thermography images (**N**) and quantitation of surface temperature over the liver (**O**) and BAT regions (**P**) in control and LKO mice (n = 4 mice). **Q**-**R**, 3D reconstruction of LD fluorescence images (**Q**) and LD size comparisons (**R**) in control and LKO hepatocytes cultured *in vitro* for 0 or 12 hours (n = 1000-4000 LDs). **S**-**T**, Respiration assay of freshly isolated primary hepatocytes from control and LKO mice. O_2_ concentration and the time derivative traces (**S**) and quantitation of O_2_ consumption rates are shown (**T**) (n = 4 biological replicates). The bracket in **S** denotes the time window during which O_2_ consumption rate was sampled. Scale bar: 100 μm in **D**, 20 μm in the inset; 3 μm in **E** and **Q**. Data are mean ± s.e.m. Statistical significance was calculated by unpaired Student’s two-sided t-test (**A**, **B**, **C**, **G**, **H**, **L**, **O**, **P**, and **T**), or two-way ANOVA with repeated measurements (**J**, **K**, and **M**), or one-way ANOVA with Tukey’s post hoc correction for p values in regular type in **R**, and one-way ANOVA with Tukey’s correction based on the sample size, means, and standard errors derived from two-way ANOVA with Tukey’s correction for p values in bold in **R**.

Multiple lines of observations suggest that impaired LD biogenesis in CLSTN3B-deficient hepatocytes results in more profound FAO. Firstly, no concomitant increases in serum cholesterol, TAG, and free fatty acid (FFA) levels, as well as adipose tissue masses, were observed (Fig. 2A, S2P-S), suggesting that the reduction in hepatic lipid accumulation was not due to enhanced lipid export or redistribution to peripheral tissues. Secondly, the LKO mice displayed increased energy expenditure as indirect calorimetry revealed significantly higher O_2_ consumption, CO_2_ production, and heat production in LKO mice on HFD for 8 weeks than the control mice, without notable differences in locomotive activity (Fig. 2J-K, S2T-U). Rectal temperature was also significantly higher in LKO mice at ZT18 (Fig. 2L). Thirdly, despite comparable food intake (Fig. S2V), the LKO mice exhibited significantly slower weight gain upon HFD feeding (Fig. 2M, two-way ANOVA, p_genotype x time_=0.018) without a significant difference in body composition (Fig. S2W), indicating reduced metabolic efficiency. Infrared thermography showed that the LKO mice exhibited higher surface temperature over the liver region but not over brown adipose tissue (BAT), which instead trended cooler (Fig. 2N-P), suggesting increased hepatic metabolic activity. At the cellular level, primary LKO adipocytes exhibited significantly faster rates of lipid storage loss when cultured *in vitro* (Fig. 2Q-R), accompanied by higher respiration rates (Fig. 2S-T). Gene expression analysis revealed that the mRNA levels of peroxisome proliferator-activated receptor gamma coactivator 1-alpha (PGC1α), a key promoter of mitochondrial biogenesis, and PPARα, the nuclear receptor regulating genes involved in FAO, along with multiple PPARα target genes involved in FAO, are upregulated in the LKO liver (Fig. S2X), whereas the FAO Blue fluorescence assays confirmed increased FAO enzymatic activity in LKO hepatocytes (Fig. S2Y-Z). Taken together, the findings show that diet-induced CLSTN3B expression in hepatocytes is crucial for supporting LD biogenesis and lipid storage, whereas CLSTN3B ablation diverts more lipids towards oxidation rather than storage, resulting in higher energy expenditure and reduced metabolic efficiency observed at the animal level.

### Transgenic overexpression of CLSTN3B promotes LD biogenesis and lipid deposition in hepatocytes

To complement the loss-of-function approach, we generated a liver-specific CLSTN3B gain-of-function mouse model to understand how CLSTN3B may influence hepatocyte LD biogenesis and lipid storage (LTG: *albumin-cre*; *Clstn3b*^tg/0^)(*38*). Even under the chow diet condition, the LTG mice exhibited significantly increased liver mass, liver-to-body weight ratio, and hepatic TAG content compared to the WT control (Fig. S3A-F). When subjected to HFD feeding, LTG mice showed significantly elevated liver-to-body weight ratio and hepatic TAG contents (Fig. 3A-C). Although few visible LDs could be identified by histological analysis (Fig. 3D), the gross appearance of the liver and the thickened buoyant LD layer from liver homogenates both indicate increased hepatic lipid deposition in LTG mice (Fig. 3D).

**Figure 3.**
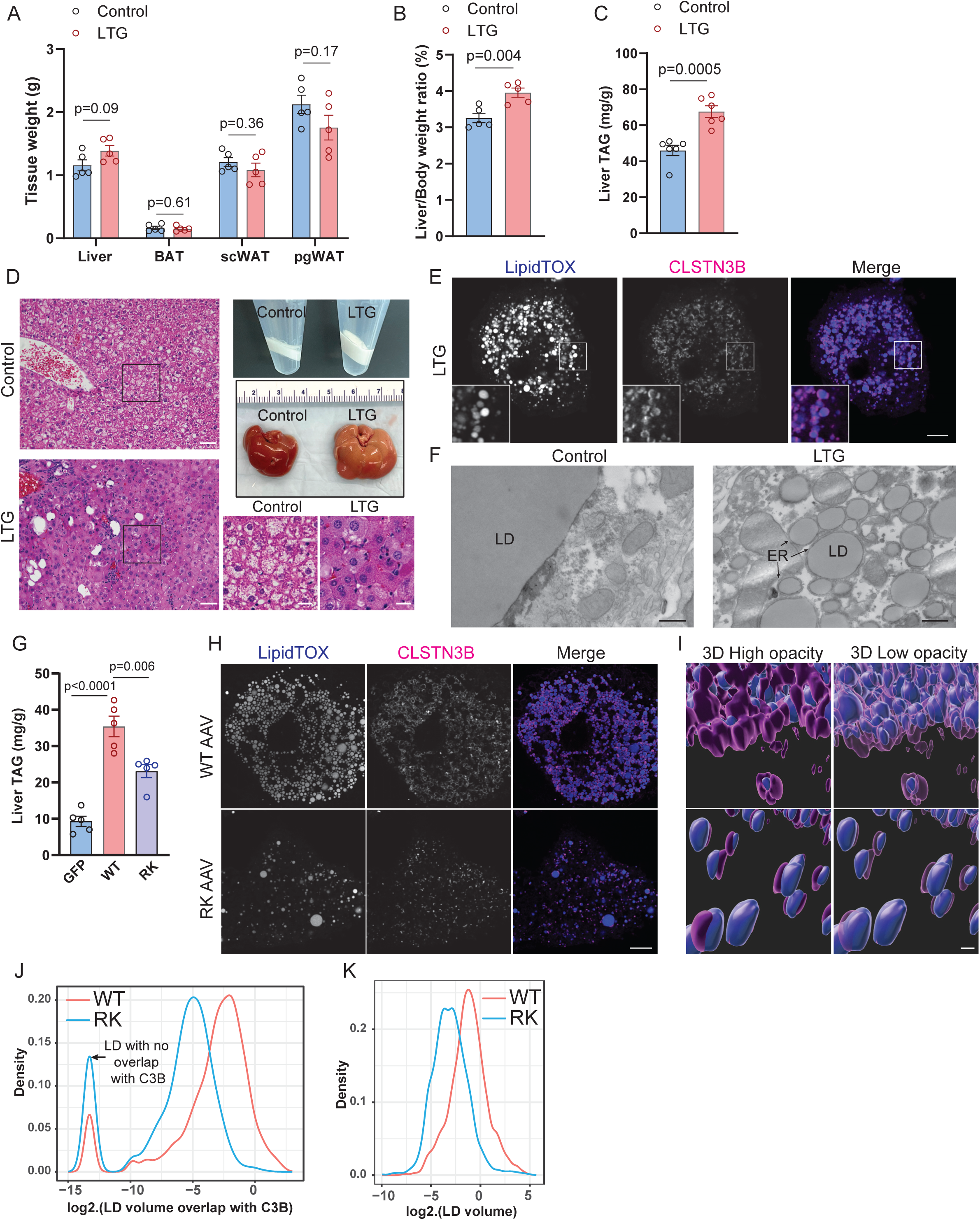
Transgenic overexpression of CLSTN3B in hepatocytes promotes lipid accumulation and exacerbates hepatic steatosis. **A**-**F**, Tissue weights (**A**), liver/body weight ratio (**B**), liver TAG content (**C**), representative gross liver images, buoyant LD fraction appearance and histology (**D**), fluorescence microscopy of primary hepatocytes (**E**), and electron microscopy of liver sections (**F**) from control and LTG mice on HFD for 10 weeks (n = 5 mice). **G**-**K**, Liver TAG content (**G**) (n = 5 mice), fluorescence microscopy of primary hepatocytes (**H**), 3D reconstruction (**l**), LD/CLSTN3B contact volume distribution curves (**J**), and LD size distribution curves (**K**) in hepatocytes from mice receiving AAV-WT-CLSTN3B or AAV-RK-CLSTN3B injection (n = 3300 for WT and 2300 for RK). CLSTN3B surface opacity is adjusted in **l** to allow visualization of the LD surface enwrapped inside the CLSTN3B surface. The arrow in **J** denotes LDs not in contact with CLSTN3B signal but assigned with a small value to permit visualization on a logarithmic scale. Scale bar: 100 μm in **D**, 20 μm in the inset; 5 μm in **E** and **H**; 500 nm in **F** and **l**. Data are mean ± s.e.m. Statistical significance was calculated by one-way ANOVA with Tukey’s post hoc test (**G**) or unpaired Student’s two-sided t-test (all other panels).

Fluorescence and electron microscopy revealed a large number of small LDs in LTG hepatocytes, characterized by robust CLSTN3B signals on the surface and extensive contact with the ER (Fig. 3E-F), mirroring previous observations made in adipocytes and other cell types(*36*) and confirming those LDs are too small to be seen by standard H&E staining. Notably, serum TAG and cholesterol levels were reduced in the LTG mice (Fig. S3G), suggesting that CLSTN3B promotes intrahepatocellular lipid storage in cytosolic LDs at the expense of lipid output via the lipoprotein pathway. Contrary to the liver LKO, the LTG liver exhibited decreased expression levels of mitochondrial FAO genes (Fig. S3H). Similar phenotypes were observed with male LTG mice and control littermates on a WD (Fig. S3I-M) or Female LTG mice on a HFD (Fig. S3N-Q).

Our observations with the LTG mice strongly support that CLSTN3B expression is sufficient to promote LD biogenesis and lipid storage in hepatocytes by inducing ER/LD hemifusion-like membrane bridge formation. To test this ideal, we used AAV to exogenously express WT CLSTN3B or the RK mutant (10 R residues replaced with K) in the livers of mice on a chow diet. Despite comparable expression levels (Fig. S3R-S), mice receiving WT CLSTN3B accumulated significantly more hepatic TAGs than those expressing the RK mutant (Fig. 3G). Fluorescence imaging with 3D reconstruction further revealed more extensive CLSTN3B localization on LD surfaces and larger LD sizes in WT-versus RK-expressing hepatocytes (Fig. 3H-K). Collectively, these data demonstrate that hepatocyte CLSTN3B expression promotes LD biogenesis and lipid storage in hepatocytes (steatosis), and its R-rich segment is essential for this effect.

### Hepatic CLSTN3B expression exacerbates MASLD pathology

We then examined whether hepatic CLSTN3B expression impacts MASLD pathology beyond simple steatosis. To this end, we subjected the LKO and LTG mice, along with their respective control mice, to a WD for 16 weeks, a regimen that more robustly induces fibrosis than HFD alone. Fluorescence imaging revealed markedly reduced α-SMA and F4/80 staining in LKO liver sections compared to controls (Fig. 4A-C), indicating attenuated hepatic fibrosis and decreased macrophage infiltration. Consistently, Sirius Red staining confirmed lower collagen deposition in LKO livers (Fig. 4A and 4D). In contrast, LTG mice displayed exacerbated fibrosis and increased macrophage infiltration relative to controls (Fig. 4A-D). Gene expression analyses further supported these histological findings: LKO livers exhibited significantly reduced expression of pro-fibrotic and pro-inflammatory genes, while LTG livers showed elevated levels of these transcripts (Fig. 4E-F). Similarly, AAV-induced CLSTN3B ablation or overexpression phenocopied the respective LKO and LTG models (Fig. S4A-D), confirming that modulating CLSTN3B expression precisely during WD feeding is sufficient to impact steatohepatitis and fibrosis. Furthermore, WD-fed LKO mice exhibited significantly lower serum aspartate aminotransferase (AST) and alanine aminotransferase (ALT) levels (Fig. 4G), consistent with alleviated liver injury, whereas substantial elevations in serum AST and ALT levels were observed in the LTG mice, indicating exacerbated liver injury (Fig. 4H).

**Figure 4.**
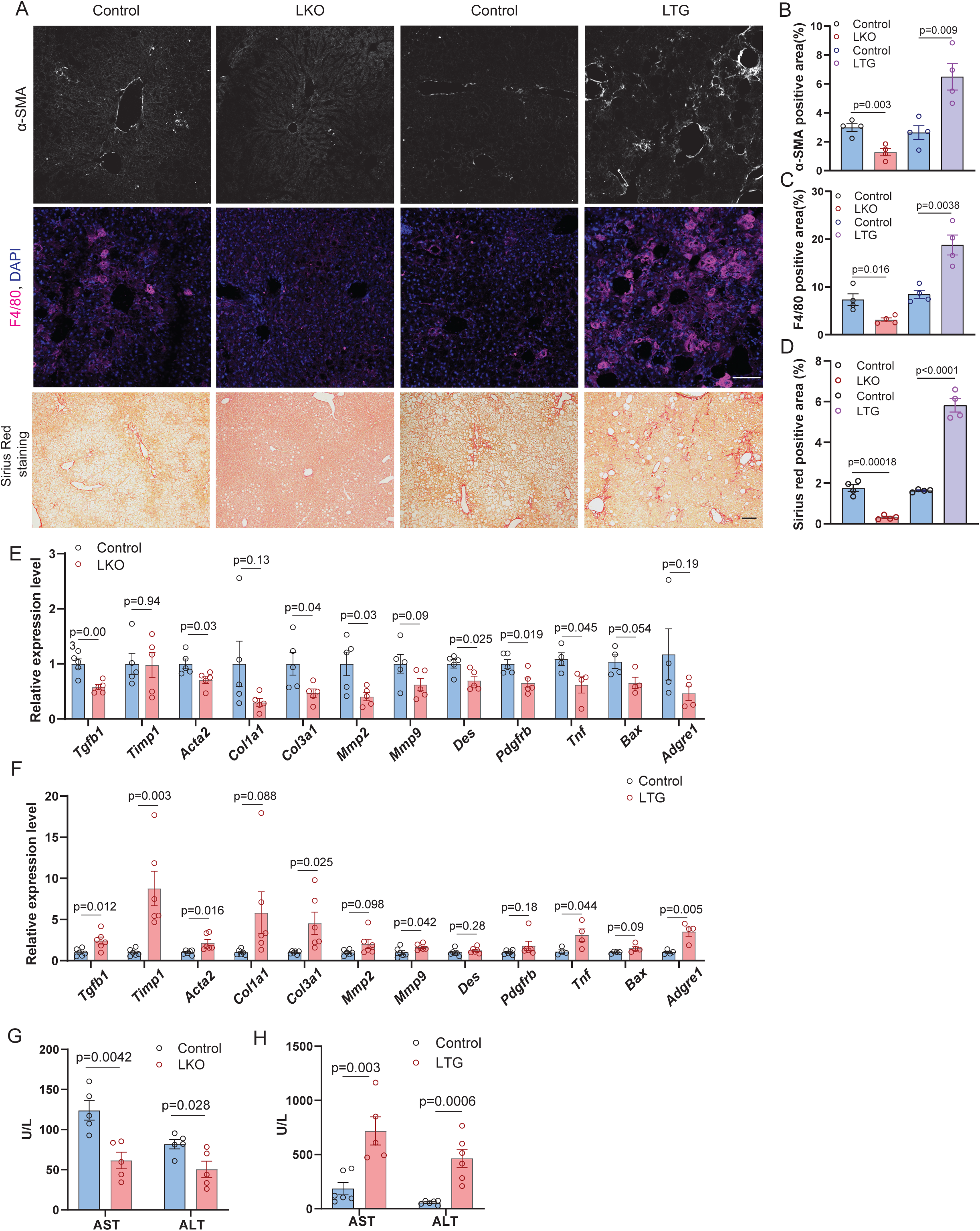
Hepatic CLSTN3B expression exacerbates MASLD pathology. **A**-**D**, Immunostaining of the fibrosis marker α-SMA, the macrophage marker F4/80, and Sirius Red staining in liver sections of control, LKO, and LTG mice (n = 4 mice) on WD for 16 weeks. Representative images (**A**) and quantitation of α-SMA- (**B**), F4/80- (**C**), and Sirius Red-positive areas (**D**) are shown. **E**-**F**, qPCR analysis of genes associated with hepatic fibrosis and inflammation in livers of control and LKO mice (**E**) or control and LTG mice (**F**) (n = 4-5 mice). **G**-**H**, Serum AST and ALT levels in control and LKO mice (**G**) or control and LTG mice (**H**) on WD for 10 weeks (n = 5-6 mice). Scale bar: 100 μm. Data are mean ± s.e.m. Statistical significance was calculated by unpaired Student’s two-sided t-test.

To determine whether MASLD pathology can be reversed by acute suppression of CLSTN3B, we delivered *Clstn3b*-targeting siRNA encapsulated in lipid nanoparticles (LNPs)(*46*) to mice on a WD. Two injections at a 0.5 mg/kg siRNA dose over three days achieved more than 90% reduction of hepatic *Clstn3b* transcript levels after 2 weeks (Fig. S5A-B). Compared with mice receiving control siRNA, mice dosed with *Clstn3b*-targeting siRNA LNPs exhibited marked improvement across all major features of MASLD, including hepatic steatosis, hepatocellular injury, and fibrosis (Fig. S5C-J). These findings identify CLSTN3B as a viable therapeutic target for reversing established MASLD pathology.

Taken together, these results demonstrate that hepatic CLSTN3B expression induces the full spectrum of MASLD pathology beyond simple steatosis. Moreover, the LTG mice represent a novel and powerful genetic model featuring rapid recapitulation of MASLD progression.

### CLSTN3B restricts mitochondrial linoleic acid availability, cardiolipin synthesis, and exacerbates oxidative stress

We next investigated the mechanistic link between CLSTN3B-driven LD biogenesis and hepatocellular injury, an unresolved yet central question in MASLD pathogenesis. A recent study demonstrated that impaired CL maturation is a hallmark of human MASLD and that a mouse model of hepatic CL deficiency recapitulates key features of MASLD pathology(*35*). Because mitochondrial CL synthesis and remodeling depend on phospholipid supplied from the ER, we hypothesized that CLSTN3B-driven LD biogenesis at the ER may disrupt normal lipid tracking and restrict mitochondrial CL production. To test this hypothesis, we performed mitochondrial and LD lipidomics analyses on livers from WT and LKO mice. Mitochondrial lipidomics revealed significantly higher levels of CLs in LKO mitochondria compared with WT controls, whereas the magnitude of the increase progressively amplified with advancing CL maturation (Fig. 5A). Consistently, analysis of CL fatty acid composition revealed an increased proportion of linoleoyl (18:2) and a decreased proportion of oleoyl (18:1) acyl chains in LKO mitochondria relative to WT controls (Fig. 5B). CL maturation depends on tafazzin-mediated transacylation using linoleoylated phosphatidylcholine (PC) and phosphatidylethanolamine (PE) as acyl donors. Indeed, LKO mitochondria exhibited significantly increased fractions of linoleoyl (18:2) acyl chains, but not oleoyl (18:1) or palmitoyl (18:0), in both PC and PE (Fig. 5C-D). In contrast, LD lipidomics revealed a significant reduction in linoleoylated TAG species in LKO hepatocytes relative to WT, accompanied by a significant increase in the fraction of saturated acyl chains (14:0 and 16:0), (for each FA, the fractional abundance in TAG in both the control and LKO LD or the relative level to control is shown in Fig. 5E). Together, these data indicate that CLSTN3B expression promotes sequestration of LA into LD TAGs, whereas CLSTN3B ablation increases the availability of linoleoylated phospholipids for mitochondrial CL maturation.

**Figure 5.**
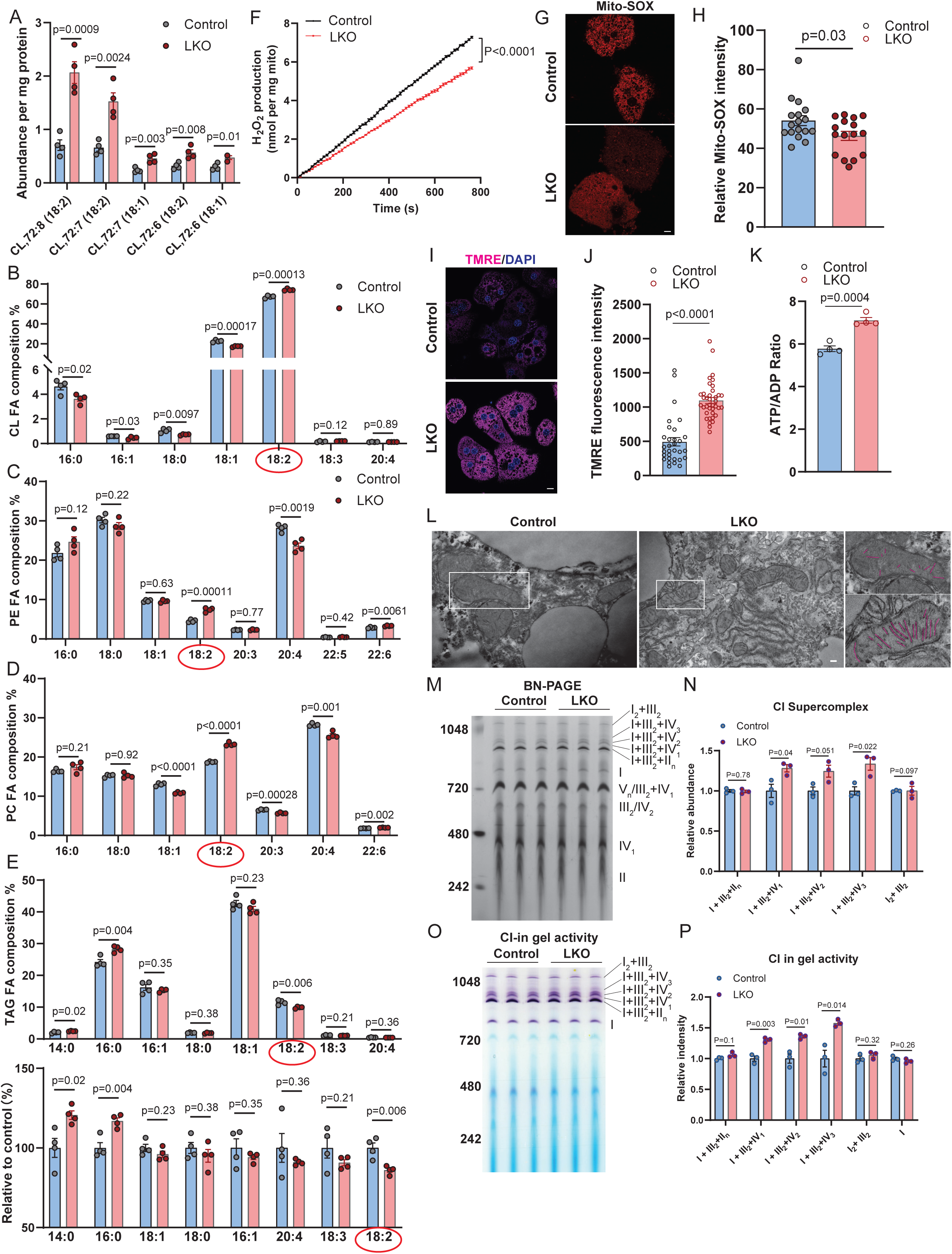
CLSTN3B restricts mitochondrial linoleic acid availability, CL synthesis, andexerbates oxidative stress. **A**-**B**, Abundance of individual CL FA species normalized to mitochondrial protein content (**A**) or expressed as fraction in total CL FA (**B**) in mitochondria isolated from control or LKO mice on WD for 10 weeks (n = 4 mice). **C**-**D**, Fractional composition of individual fatty acyl chains in mitochondrial PE (**C**) and PC (**D**) from control or LKO mice on HFD for 10 weeks (n = 4 mice). **E**, Fractional composition of individual fatty acyl chains in LD TAGs from control or LKO mice on WD for 10 weeks (n = 4 mice). The FAs are arranged in a decreasing order in the lower plot showing the relative levels to control. **F**. Time course of electron leak (H_2_O_2_ production) from mitochondria isolated from control or LKO liver respiring on succinate (n = 4 biological replicates). **G**-**H**, Representative images (**G**) and quantitation (**H**) of mito-SOX staining in control or LKO primary hepatocytes (n = 16-17 cells). **I**-**J**, Representative images (**I**) and quantitation (**J**) of TMRE staining in control or LKO primary hepatocytes (n = 31-38 cells). **K**, Quantitation of ATP/ADP ratio in primary control and LKO hepatocytes (n = 4 biological replicates). **L**, Representative electron microscopic images of liver sections from control and LKO from mice on WD for 10 weeks. Cristae architecture is outlined in magenta. **M**-**N**, Blue native PAGE analysis (**M**) and quantitation (**N**) of ETC supercomplexes from control and LKO liver (n = 3 mice). **O**-**P**, Complex I in-gel activity assay (**O**) and quantitation (**P**) of complex I-containing ETC supercomplexes from control and LKO liver (n = 3 mice). Scale bar: 5 μm in **G** and **I**; 200 nm in **L**. Data are mean ± s.e.m. Statistical significance was calculated by two-way ANOVA with repeated measurements (**F**) or unpaired Student’s two-sided t-test (all other panels).

Given that CL deficiency is known to increase electron leak from the ETC, exacerbating oxidative stress and hepatocyte injury(*35*), we next assessed mitochondrial function. Mitochondria isolated from LKO hepatocytes exhibited significantly reduced electron leak, as measured by H_2_O_2_ production during succinate-driven respiration (Fig. 5F). Consistently, LKO hepatocytes displayed reduced mitochondrial ROS levels by MitoSOX staining (Fig. 5G-H), increased mitochondrial membrane potential as assessed by tetramethylrhodamine ethyl ester (TMRE) staining (Fig. 5I-J), and an elevated ATP/ADP ratio compared with WT hepatocytes (Fig. 5K).

We further examined mitochondrial ultrastructure and ETC organization. Transmission electron microscopy revealed disrupted cristae morphology and irregular cristae packing in WT hepatocytes relative to LKO hepatocytes (Fig. 5L). Blue-native PAGE followed by Coomassie staining and in-gel activity assays showed reduced ETC supercomplex formation in WT hepatocytes compared with the LKO, particularly the I+III_2_+IV_n_ supercomplexes (Fig. 5M-P). This pattern mirrors ETC defects observed in Barth Syndrome patients harboring tafazzin mutations, who are unable to generate mature CL(*47*), and mice with hepatic deficiency of cardiolipin synthase(*35*).

Collectively, these findings demonstrate that CLSTN3B ablation enhances mitochondrial CL maturation by increasing the availability of linoleoylated phospholipids, thereby improving mitochondrial bioenergetic efficiency, reducing oxidative stress, and preserving hepatocyte integrity, even under conditions of high FAO-derived electron flux. This mechanism provides a direct link between ER-driven LD biogenesis, mitochondrial phospholipid remodeling, and hepatocellular injury, offering a unifying explanation for the protective effects of CLSTN3B deficiency against MASLD progression.

### Linoleic acid facilitates CLSTN3B-driven LD biogenesis

We next sought to determine why CLSTN3B-driven LD biogenesis preferentially sequesters LA into TAG stored within LDs. A central aspect of CLSTN3B activity is that it interacts with the negatively charged, cone-shaped phosphatidic acid (PA) to support the hemifusion-like membrane bridge at ER/LD junction via its R-rich region, which was demonstrated by our previously published liposome fusion assay(*36*) and a lipid strip-based binding assay (Fig. S6A). Compared with saturated palmitoyl (18:0) or monounsaturated oleoyl (18:1) PA, PA with di-unsaturated linoleoyl acyl chain possesses increased conformational flexibility and a larger cross-sectional area, features that further enhance its cone geometry. These properties are ideally suited to stabilize the high negative membrane curvature characteristic of the hemifusion-like ER-LD interface. We therefore hypothesized that the unique biophysical properties of the linoleoyl acyl chain promote enrichment of LA-containing PA at CLSTN3B-supported ER/LD membrane bridges, where it can be efficiently converted into LA-containing DAG and TAG. Consistent with this model, lipidomic analysis revealed higher levels of LA-containing PA and DAG species in LDs isolated from WT compared with LKO hepatocytes (Fig. 6A-B). In contrast, LA enrichment was not observed for PC and PE in the LD fraction, again supporting the unique preference of CLSTN3B for PA (Fig. S6B-C).

**Figure 6.**
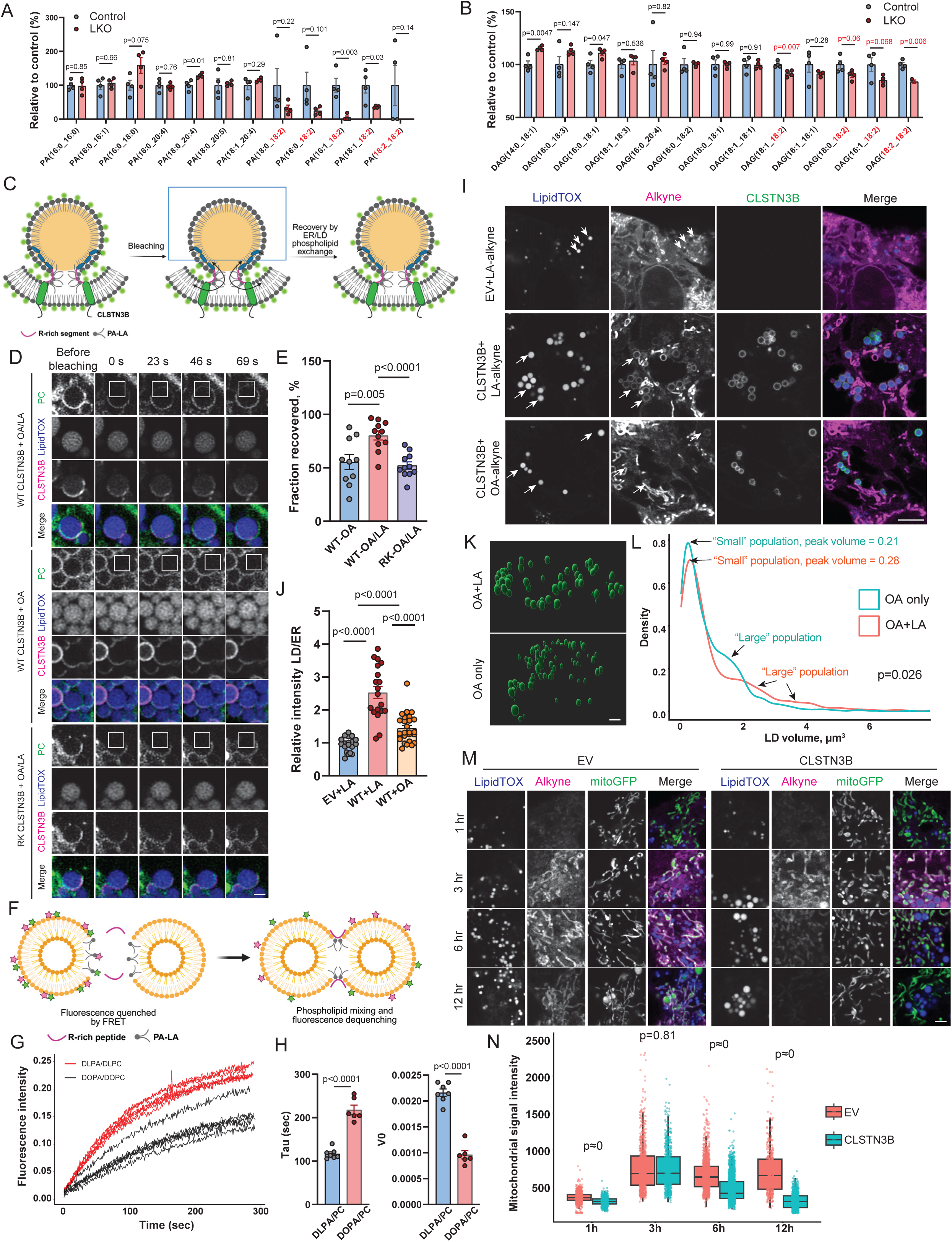
Linoleic acid facilitates CLSTN3B-driven LD biogenesis. **A**-**B**, Relative abundance of individual PA (**A**) or DAG (**B**) species to control levels in LD fractions isolated from control or LKO mice on WD for 10 weeks (n = 4 mice). **C**-**E**, Schematic overview (**C**), representative FRAP images (**D**), and quantitation (**E**) of phospholipid exchange between ER and LD using copper-free click-chemistry labeled PC (n = 10-11 LDs for each condition). **F**-**H**, Schematic overview (**F**), FRET traces (**G**), and kinetics quantitation (**H**) of CLSTN3B-derived R-rich peptide-induced lipid mixing between DLPA/DLPC or DOPA/DOPC-based liposomes (n = 6-7 replicates). **I**-**J**, Representative fluorescence images (**I**) of cells labeled with LA- or OA-alkyne and quantitation of fluorescence intensity on LDs (**J**) (n = 19-26 cells). **K**-**L**, Representative 3D reconstruction LD images (**K**) and density curves of LD size distribution (**L**) in cells treated with OA+LA or OA alone (n = 900-1200 LDs). **M**-**N**, Representative fluorescence images of cells labeled with LA-alkyne +/- CLSTN3B expression (**M**) and quantitation of mitochondrial fluorescence intensity (**N**) (n = 400-1900 mitochondria per condition). Scale bar: 1 μm in **D**; 5 μm in **I** and **M**; and 2 μm in **K**. Data are mean ± s.e.m. Statistical significance was calculated by one-way ANOVA with Tukey post hoc correction (**E** and **J**), or unpaired Student’s two-sided t-test (**H**), or Wilcoxon rank-sum test (**L** and **N**).

To directly assess whether LA-containing PA preferentially supports CLSTN3B-mediated membrane fusion, we employed complementary in-cell imaging and *in vitro* reconstitution approaches. For the imaging study, we combined copper-free click chemistry labeling of newly synthesized PC with fluorescence recovery after photobleaching (FRAP) to quantify ER-LD phospholipid exchange dynamics. We hypothesized that LA-dependent stabilization of CLSTN3B-mediated membrane bridges would enhance ER/LD phospholipid exchange, manifesting as accelerated recovery of labeled LD surface phospholipids (Fig. 6C). To test this, U2OS cells were metabolically labeled with azide-choline, induced to form LDs with either OA plus LA (50 μM each) or OA alone (100 μM), and transfected with CLSTN3B-mCherry or an RK mutant compromised in PA binding. Cells treated with OA+LA exhibited significantly faster and more extensive recovery of LD monolayer PC fluorescence following photobleaching compared with OA-treated cells (Fig. 6D-E). This recovery was markedly attenuated in cells expressing the RK mutant (Fig. 6D-E), indicating that the R-rich segment/PA-LA interaction-supported ER-LD membrane bridging enhances phospholipid exchange.

To further interrogate this mechanism in a reconstituted system, we performed an *in vitro* FRET-based liposome fusion/lipid-mixing assay using liposomes containing either di-linoleoyl (DLPA:DPLC = 6:4) or di-oleoyl phospholipids (DOPA:DOPC = 6:4) (Fig. 6F). The R-rich peptide of CLSTN3B induced significantly faster lipid mixing in liposomes containing di-linoleoyl PA/PC than in those containing di-oleoyl PA/PC (Fig. 6G-H), supporting the notion that linoleoyl acyl chains lower the energy of hemifusion intermediates and thus are likely to be favored at CLSTN3B-supported hemifusion-like membrane bridges with negative curvature.

The enrichment of LA-containing PA on LDs, together with its enhanced ability to support CLSTN3B-driven membrane bridging, predicts that LA should selectively promote CLSTN3B-dependent LD biogenesis. To test this, we examined LA partitioning in HEK293 cells exogenously expressing CLSTN3B using an alkyne derivative of LA combined with click-chemistry imaging. Strikingly, CLSTN3B expression led to pronounced enrichment of LA on the LD surface compared with control cells (Fig. 6I-J). In contrast, the alkyne derivative of monounsaturated oleic acid exhibited substantially diminished LD surface enrichment, despite CLSTN3B expression (Fig. 6I-J). These findings demonstrate that CLSTN3B selectively directs LA to the ER-LD interface across cellular contexts beyond hepatocytes. Furthermore, CLSTN3B-expressing cells exhibited more rapid LD expansion when supplied with OA+LA than with OA alone (Fig. S6D-E) and eventually reached significantly larger LD volumes (Figure 6K-L).

Notably, LA-alkyne labeling was also prominent on mitochondria (Fig. 6M), indicating trafficking of LA-containing phospholipids to mitochondria. We therefore compared mitochondrial LA labeling in cells with or without CLSTN3B expression. In cells lacking CLSTN3B expression, mitochondrial LA labeling remained relatively stable over the 3-12 hr time window. In contrast, CLSTN3B-expressing cells exhibited a progressive reduction in mitochondrial LA labeling concomitant with increases in total LD volume during this period (Fig. 6M-N). These observations are consistent with diversion of LA away from mitochondria in steatotic hepatocytes.

Taken together, these data support a model in which CLSTN3B selectively directs PA-LA to ER-LD membrane bridges, where they are efficiently converted into TAG and sequestered within LDs, thereby limiting LA availability to mitochondria. This interorganelle competition for LA provides a mechanistic explanation for how excessive CLSTN3B-driven LD biogenesis compromises mitochondrial CL maturation, elevates ROS production, and promotes oxidative stress.

### Hepatic *CLSTN3B* expression level positively correlates with clinical MASLD fibrosis severity and progression

Lastly, we sought to determine the clinical relevance of hepatic *CLSTN3B* expression in MASLD, which is characterized by progressive liver fibrosis due to chronic steatohepatitis, in clinical cohorts of MASLD patients. First, we developed a method to accurately quantitate *CLSTN3B* expression without interference from the *CLSTN3* gene. This is particularly important for analyzing human data because *CLSTN3* is expressed at a substantially higher level than *CLSTN3B* in human liver. Per the GENCODE annotation release 45, *CLSTN3B* is defined as one of the *CLSTN3* transcripts (ENST00000535313.2). *CLSTN3B* comprises three exons, with the first exon being unique. The second and third exons are shared with *CLSTN3*. To differentiate *CLSTN3B* expression from *CLSTN3*, we removed two shared exons (chr12: 7,157,489-7,157,691, chr12: 7,157,941-7,158,305) from either gene records. This left only the first exon of *CLSTN3B* (chr12: 7,155,868-7,156,929) and the remaining exons of *CLSTN3* to allow us to separately and accurately quantitate *CLSTN3B* and *CLSTN3* expression.

With this method, we found that both CLSTN3B and CLSTN3 expression were higher in 206 European MASLD patients compared to 10 healthy obese controls (p=0.000068 and p=0.0017, respectively) (Fig. 7A, Fig. S7A). Among the MASLD patients, expression levels of CLSTN3B, but not CLSTN3, were higher in patients with advanced liver fibrosis or cirrhosis (i.e., histological fibrosis stage F3-4) compared to the rest of the patients with no to modest fibrosis (i.e., F0-2) (p=0.046) (Fig. 7B, Fig. S7B). The association was replicated in the two cohorts of MASLD patients in the U.S. (n=78, p=0.037; n=112, p=0.039) (Fig. 7C and D), and another cohort of 106 non-cirrhotic Japanese MASLD patients (p=0.03) (Supplementary Table 2). In 77 patients from the Japanese cohort with paired liver biopsy tissues obtained at a median interval of 2.3 years, time-adjusted progression of histological liver fibrosis stage was positively correlated with elevated baseline expression levels of CLSTN3B, but not CLSTN3 (Spearman’s rho=0.26, p=0.023) (Fig. 7E, Fig. S7C). Notably, in this Japanese cohort, a non-significant trend (p=0.055, Fisher test) was observed in the correlation between steatosis severity and CLSTN3B expression level when the comparison was made between the bottom quartile (Q1) of CLSTN3B expression level and the rest (Q2-Q4), suggesting a potential impact of CLSTN3B expression on steatosis in MASLD patients. The hepatic CLSTN3B expression was positively correlated with CLSTN3 expression (Fig. S7D, Spearman’s rho=0.29, p=0.023), but the association with fibrosis progression is unique to CLSTN3B and not affected by CLSTN3 (Supplementary Table 3).

**Figure 7.**
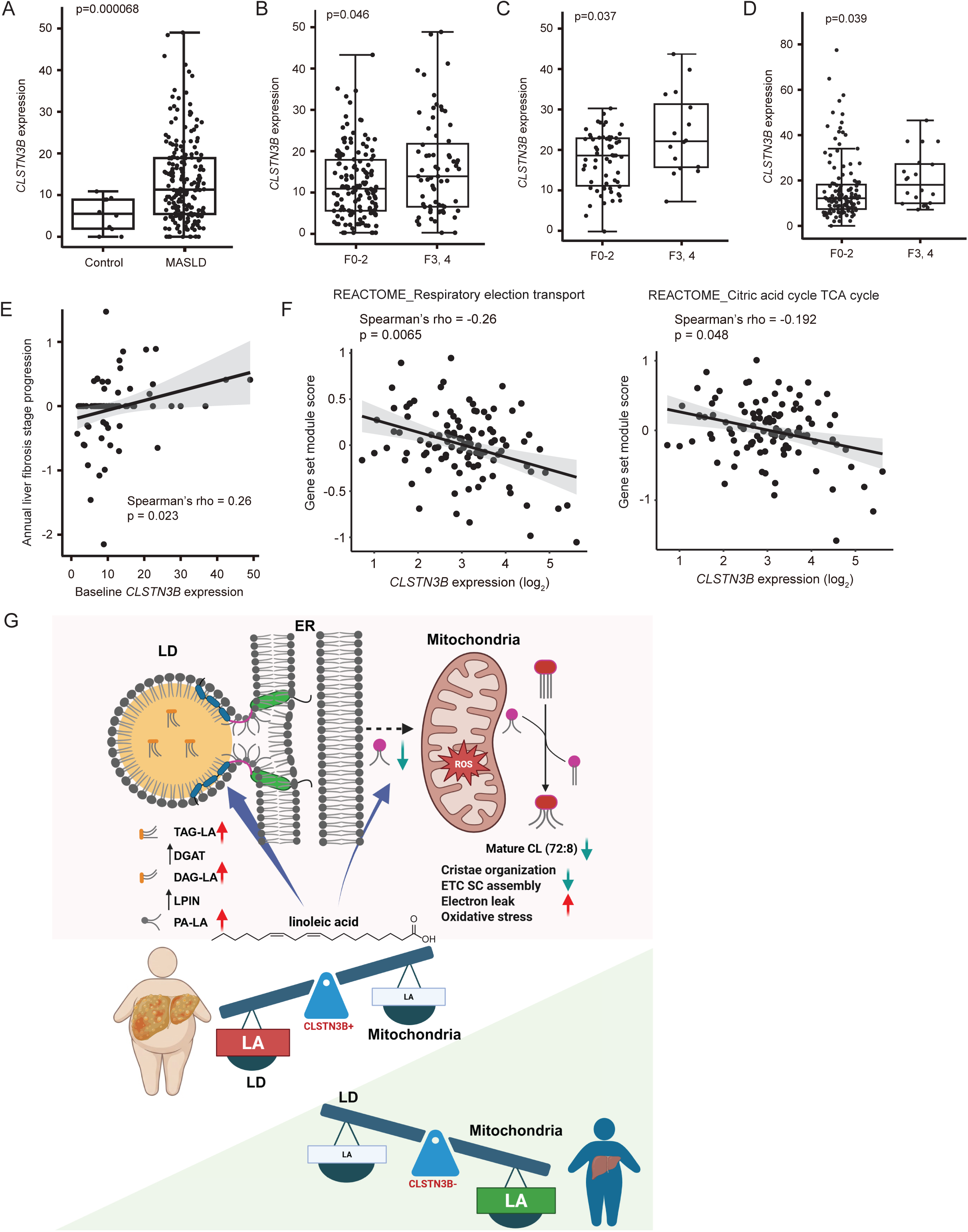
Hepatic *CLSTN3B* expression level positively correlates with clinical MASLD progression. **A**, Hepatic *CLSTN3B* expression levels in 206 European MASLD patients compared to 10 healthy obese controls. **B**-**D**, Hepatic CLSTN3B expression stratified by liver fibrosis stage (F0-2 vs. F3-4) in the European MASLD cohorts (n = 206) (**B**) and two U.S.-based MASLD cohorts (n = 78) (**C**) and (n = 112) (**D**). **E**, Correlation between baseline *CLSTN3B* expression levels and time-adjusted (i.e., annual) progression of liver fibrosis stage in 77 Japanese MASLD patients. Inter-group difference was assessed by Wilcoxon rank-sum test. Correlation was assessed by Spearman correlation test. **F**, Correlation between OXPHOS- TCA cycle-related genes with CLSTN3B expression levels in the Japanase MASLD cohort (n = 106). **G**, Model of CLSTN3B’s effect on interorganelle competition for linoleic acid in hepatocytes.

Lastly, we explored molecular pathways associated with CLSTN3B expression in transcriptome profiles in the Japanese MASLD cohort. Among the genes involved in the PPARγ/RXR/CEBP transcriptional axis, the key adipogenic regulators, CEBPA showed significant positive correlations with CLSTN3B expression (Speaman’s rho=0.39, p= 0.000031; Supplementary Table 4). In contrast, CLSTN3B expression was significantly negatively correlated with gene sets related to oxidative phosphorylation (OXPHOS) and the tricarboxylic acid (TCA) cycle (Fig. 7F), further supporting a role for CLSTN3B in promoting lipid storage while concomitantly suppressing mitochondrial activity (Fig. 7G).

## Discussion

Our study identifies CLSTN3B as a previously unrecognized regulator of LD biogenesis and systemic energy homeostasis in hepatocytes, whose expression is robustly induced by dietary caloric overload through PPARγ activation. Using complementary gain- and loss-of-function mouse models, we demonstrate that CLSTN3B promotes LD biogenesis and hepatic lipid storage by facilitating the formation of ER/LD membrane bridges via its R-rich segment. This activity selectively enriches PA with linoleoyl acyl chains at ER/LD junctions and drives preferential sequestration of LA into LD TAGs. As a consequence, mitochondrial LA availability is reduced, leading to impaired CL synthesis, defective ETC supercomplex assembly, increased electron leak, oxidative stress, and exacerbated hepatocyte injury (Figure 7G). Consistent with this model, CLSTN3B ablation markedly impairs LD biogenesis and ameliorates all major pathological features of MASLD. These findings align with previous studies showing that deficiencies in LD-associated proteins such as PLIN2 and CIDEB impair LD biogenesis and lipid storage while mitigating MASLD pathology(*25–30*). An intriguing implication of our work is that reduced LD biogenesis in the PLIN2 or CIDEB models may also restore mitochondrial LA and CL levels, a possibility that warrants direct experimental testing. Taken together, our results provide a mechanistic explanation for how excessive LD biogenesis in hepatocytes drives cellular injury and disease progression. In this framework, CLSTN3B induction represents an adaptive response whereby hepatocytes recruit an adipocyte-selective LD biogenesis program to accommodate excess lipid influx. However, this adaptation becomes maladaptive by disrupting normal interorganelle lipid partitioning and depleting mitochondrial LA, thereby compromising mitochondrial integrity and function.

Our findings in hepatocytes reinforce prior conclusions from the adipocyte system, underscoring the core function of CLSTN3B as a key regulator of LD biogenesis by facilitating lipid and protein trafficking via the hemifusion-like membrane bridge between ER and LD stabilized by its arginine-rich segment(*36*). This may seem paradoxical, as CLSTN3B is most abundantly expressed in brown adipocytes, which are better known for FAO and thermogenesis rather than lipid storage(*37, 38*). However, LD maturation is critical for shaping the lipolytic response of brown adipocytes: immature LDs exhibit elevated basal lipolysis but reduced responsiveness to hormonal stimulation(*36*). Moreover, CLSTN3B promotes sympathetic innervation of brown adipocytes via an S100B-dependent mechanism(*38*), a function not observed in white adipocytes or hepatocytes, which receive minimal direct sympathetic input. Therefore, in white adipocytes and hepatocytes, where neural stimulation of lipid mobilization is limited, the primary manifestation of CLSTN3B-mediated LD maturation appears to be enhancement of lipid storage capacity.

A recent study reported that CLSTN3B is upregulated in mouse hepatocytes in a PPARα-dependent manner during ketogenic diet (KD) feeding and is required for efficient FAO and ketogenesis(*44*). This observation seems at odds with our finding that CLSTN3B promotes lipid storage under HFD or Western diet WD conditions. We propose that this discrepancy may reflect another case of context-dependent outcomes of LD maturation. Under ketogenic conditions, CLSTN3B may facilitate efficient fatty acid channeling from LD toward mitochondria, whereas under HFD or WD conditions, it instead promotes lipid sequestration and mitochondrial lipid deprivation. Resolving this apparent contradiction will require systematic comparisons of LD composition, interorganelle communication, and lipid flux across distinct dietary states.

Our observed preferential sequestration of LA into hepatic TAGs during steatosis aligns with a previous study showing that LA is enriched in the liver of a mouse MASLD model(*48*). Importantly, we provide a mechanistic explanation for this phenomenon from a biophysical perspective. The di-unsaturated linoleoyl acyl chain confers enhanced conformational flexibility and conical geometry, favoring stabilization of the high negative curvature characteristic of hemifusion-like ER/LD membrane bridges. This principle may extend beyond hepatocytes to other tissues in which pathological LD accumulation coincides with mitochondrial dysfunction. For example, cardiomyocyte-specific PPARγ transgenic models of lipotoxic cardiomyopathy exhibit disrupted mitochondrial cristae architecture alongside lipid accumulation(*7*). Given that tetralinoleoyl CL is the dominant CL species in mammalian cardiomyocytes and is essential for cristae organization and respiration(*49*), it is tempting to speculate that excessive LD biogenesis in cardiomyocytes similarly diverts LA away from mitochondria, resulting in CL deficiency. LD accumulation has also been reported in neurons under neurodegenerative conditions(*5*). Neuronal membranes are uniquely enriched in PUFAs, particularly docosahexaenoic acid, which is critical for membrane fluidity and neuronal function. Excessive LD biogenesis in neurons could therefore lead to selective sequestration of these PUFAs into LD TAGs, depleting them from other membrane compartments and contributing to neurodegenerative pathology, a hypothesis that merits future investigation. Emerging evidence suggests potential links between dysregulated lipid metabolism, α-synuclein aggregation, and mitochondrial defects in Parkinson’s disease(*50*). Intriguingly, α-synuclein is known to bind anionic PUFA-containing phospholipids(*51, 52*), raising a more generalized possibility that protein/lipid co-aggregates, akin to LDs, may act as a sink that competes with other organelles for essential lipid species.

While our data establish CLSTN3B as a critical driver of LD biogenesis in mouse models of MASLD, other proteins involved in ER/LD junction formation may predominate in different pathological settings. We speculate that a shared feature among LD biogenesis drivers is the presence of polycationic motifs positioned between membranes. These motifs are well suited to interact with PA species containing PUFA acyl chains and to stabilize the negative curvature of hemifusion-like membrane bridges, thus effectively competing with other organelles for PUFAs. SEIPIN, a canonical LD biogenesis factor, contains a polycationic region on the cytosolic side of its ER transmembrane domain (RARR aa23-26, and RHR, aa265-267) and has been shown to bind PA(*53*). Together with its adipocyte-selective enhancer adipogenin(*54*), SEIPIN or its mutant may drive pathogenic LD biogenesis or protein/lipid aggregation, especially given its expression in the CNS and the association of SEIPIN mutations with neuropathy(*55*).

Although the molecular identity of the LD biogenesis driver may vary across disease contexts, our conditional CLSTN3B transgenic mouse model provides a powerful system to interrogate the consequences of excessive LD biogenesis and PUFA sequestration in specific cell types of interest. Its power stems from cleanly isolating lipid-driven pathogenesis and directly testing the principle that disrupting interorganelle lipid flow is sufficient to induce organelle failure and pathology. This model will be valuable for establishing a generalizable experimental paradigm for determining whether other disease contexts operate through a similar mechanism of interorganelle lipid competition, while also enabling identifying therapeutic strategies aimed at restoring balanced lipid partitioning among organelles and mitigating cellular damage arising from pathological lipid storage.

## Methods

### Mouse strains

All animal studies were approved by and in full compliance with the ethical regulation of the Institutional Animal Care and Use Committee (IACUC) of University of Texas Southwestern Medical Center. Mice were euthanized by CO_2_ asphyxiation, according to the guidelines of IACUC and the recommendations of the Panel on Euthanasia of the American Veterinary Association.

The *Clstn3b flox/flox* and the *Clstn3b* transgene mice were described in a previous publication (Zhang et al, PMID: 38293096). Mice were maintained on a normal chow diet (13% fat, 57% carbohydrate and 30% protein, PicoLab Rodent Diet 5L0D) or placed on a high-fat diet (HFD, Rodent diet with 60 kcal% Fat, Research diet, D12492, USA) or western diet consisted of high sugar, high fat, and high cholesterol solid food (Teklad Diets #TD. 120528) combined with high-sugar water containing 23.1 g/L d-fructose (Sigma-Aldrich #G8270) and 18.9 g/L d-glucose (Sigma-Aldrich #F0127) or CDA-HFD consisted of a methionine and choline-deficient high-fat diet (Research Diets #A06071302) from 6-8 weeks of age for 8 to 16 weeks. All mice were maintained under a 12 hr light/12 hr dark cycle at constant temperature (23°C or 30°C as specified) and 40–60% humidity with free access to food and water. Liver specific Clstn3b knockout mice were generated via crossing *Clstn3b flox/flox* mice with *Albcre* mice. Liver specific Clstn3b transgenic mice were generated via crossing *Clstn3b* transgene mice with *Albcre* mice. All animal studies were approved by and in full compliance with the ethical regulation of the Institutional Animal Care and Use Committee of University of Texas Southwestern Medical Center. Sample size was chosen based on literature and pilot experiment results to ensure that statistical significance could be reached. Randomization was not performed because mice were grouped based on genotype. Littermates were used for all the experiments involving the conditional KO and transgenic mice.

### AAV production and purification

AAV production and purification were described in a previous publication (Wang et al, PMID: 37040760). Briefly, AAV8 was produced using AAV-Pro 293T cells (Takara #632273) cultured in one or more 15cm dishes. Cells were plated one day before transfection at 50% confluence, which would allow the cells to reach 80-90% confluence the next day. For transfection of one 15cm dish, 10mg MOSAICS vector, 10mg pAAV2/8 (Addgene #112864), and 20mg pAdDeltaF6 (Addgene #112867) plasmids were mixed with 1ml Opti-MEM medium in one tube. In another tube, 160ml PEI solution (1mg/ml in water, pH7.0, powder from ChemCruz #sc-360988) was mixed with 1ml Opti-MEM medium. The solutions from both tubes were then mixed and incubated for 10min before adding to cell culture. 48h after transfection, the cells were scraped off the dish and collected by centrifugation at 500g for 10min. The supernatant was disinfected and discarded, and the cell pellets were lysed in 1.5ml/15cm dish lysis buffer (PBS supplemented with NaCl powder to final concentration of 200mM, and with CHAPS powder to final concentration of 0.5% (w/v)). The cell suspension was put on ice for 10min with intermittent vortexing, and then centrifuged at 20,000g for 10min at 4C. The supernatant containing the AAV was collected. To set up the gravity column for AAV purification, 0.5ml of AAV8-binding slurry beads (ThermoFisher #A30789), or enough to purify AAV from up to eight 15cm dishes, was loaded into an empty column (Bio-Rad#731-1550). After the beads were tightly packed at the bottom, they were washed with 5ml of wash buffer (PBS supplemented with NaCl powder to a final concentration of 500mM). The supernatant containing AAV was then loaded onto the column. After all of the supernatant flowed through, the beads were washed with 10ml wash buffer twice. The AAV was then eluted with 3ml elution buffer (100mM glycine, 500mM NaCl in water, pH 2.5) and the eluate was immediately neutralized with 0.12ml 1M Tris-HCl (pH 7.5-8.0). The AAV was concentrated by centrifugation at 2000g for 3-5min at 4C using an 100k Amicon Ultra Centrifugal Filter Unit (Millipore #UFC810024). After centrifugation, the volume of AAV should be equal to or less than 0.5ml. The concentrated AAV was diluted with 4-5ml AAV dialysis buffer (PBS supplemented with powders to final concentrations of 212mM NaCl and 5% sorbitol (w/v)) and centrifuged at 2000g for 3-5min at 4C. The dilution and centrifugation processes were repeated 3 times. The final concentrated AAV was transferred into a 1.5ml tube and centrifuged at 20,000g for 5min to remove debris. The supernatant was aliquoted, flash frozen using liquid nitrogen, and stored at -80C.

### AAV injection

Six-week-old male mice were anaesthetized with isoflurane. About 0.5 × 10^11^ AAV particles were delivered into mice liver via orbital injection. Mice were allowed to recover for 2-3 weeks and then treated with HFD/WD for appropriate duration. AAV2/8 vectors were described as in Wang et al, PMID: 37040760.

### AAV induced liver specific *Pparg* KO mice

Liver-specific *Pparg* knockout mice were generated using an adeno-associated virus-based CRISPR system. AAV2/8 particles encoding by MOSAICS system, U6-driven single-guide RNA (sgRNA) targeting the mouse *Pparg* gene were constructed and packaged following above instructions. The sgRNA sequence (ATAAATAAGCTTCAATCGGA) was selected from a validated mouse gRNA sequences database (dbGuide database. https://frederick.cancer.gov/downloads/mm_guide_info.csv.gz). Eight-week-old Cas9 knock-in mice (Jackson#029415) were injected via orbital vein with 5 × 10^10^ viral genomes (vg) of AAV2/8-sg*Pparg* in 100 µL sterile PBS. Control mice received AAV2/8 expressing a non-targeting sgRNA. Mice were maintained for 2-3 weeks post-injection to allow for gene editing.

### H&E, immunofluorescence (IF), and Sirius Red staining

Liver pieces were fixed in buffered formalin (Fisherbrand #245-685) for 24h with gentle shaking at 4C and then transferred to 70%EtOH for another 24h with shaking at 4C. Paraffin embedding, liver sectioning (4mm thickness), and H&E staining were performed at the UT Southwestern Tissue Management Shared Resource Core. For Cryo-section IF staining, Tissue was collected immediately after euthanizing the mice and fixed with 4% paraformaldehyde overnight. The tissue was then washed with PBS 5 times, for 10 min each time and incubated in PBS with 30% sucrose for 8 h and then frozen in Tissue-Tek O.C.T. Compound (Sakura Finetek,4583). Frozen tissue was cut into 30-μm sections on a Leica CM3050 S cryostat. Sections were briefly rinsed with PBS and blocked with PBS/0.3% Triton X-100/5% FBS overnight. Sections were then stained with αSMA antibody (14395-1-AP, Proteintech, 1:100) or F4/80 antibody (28463-1-AP, Proteintech, 1:100) for 12 hours/overnight. Sections were washed with PBS/0.03% Triton X-100/5% FBS 5 times, for 1 h each time and then stained with anti-rabbit Alexa Fluor 488 (Thermo Fisher, A-11011, 1:500) for 2 hours. Sections were washed with PBS/0.3% Triton X-100/5% FBS 3 times, for 30 min each time and stained with BODYPI and DAPI for 30 min, wash with PBS/0.3% Triton X-100/5% FBS 3 times for 10 min each time and then mounted in ProLong Diamond Antifade Mountant (Thermo Fisher, P36965).

Sirius Red staining was performed on paraffin embedded liver sections using the Picro Sirius Red Staining Kit (Abcam #ab150681) according to the manufacturer’s protocol. Images were then taken on a ZEISS LSM 900 confocal microscope. ImageJ was used to quantify IF staining.

### Serum and liver metabolic assays

At the study endpoints, blood was collected immediately after sacrificing the mouse. Blood was let standing for 30 min and then centrifuged at 3000 g for 15 minutes at 4 °C. The supernatant (serum) was analyzed for AST, ALT, cholesterol, and TAG using specific reagent kits (VITROS 8433815, 1655281, 1669829, 1336544) on a fully automated OCD Vitros 350 dry chemistry analyzer. All analyses followed the protocols provided by the reagent kit manufacturer (Ortho Clinical Diagnostics, Raritan, NJ) at the UT Southwestern Metabolic Phenotyping Core. Or samples analyzed for AST, ALT, cholesterol, FFA and TAG using specific reagent kits (Cayman 701640, 700260, 10007640, 700310, 10010303) following the protocols provided by Cayman.

### Electron microscopy

For transmission electron microscopy (TEM), mice were first perfused transcardially with 15 mL of ice-cold fixative buffer (4% paraformaldehyde, 2.5% glutaraldehyde plus 0.2% picricc acid in 0.01M phosphate buffer) under deep anesthesia. Immediately after perfusion, small liver tissue blocks (∼1 mm³) were dissected and immersed in the same fixative buffer. The tissue was then post-fixed in 1% osmium tetroxide with 1.5 % K_3_[Fe(CN)_6_] in 0.1 M sodium cacodylate buffer for 1 h at room temperature, rinsed with water and *en bloc* stained with 0.5% aqueous uranyl acetate in 25% methanol overnight at 4 degrees C. After five rinses with water, specimens were stained with 0.02M lead nitrate in 0.03M L-aspartate for 30 minutes at 60 degrees C. Samples were dehydrated with increasing concentration of ethanol and infiltrated with Embed-812 resin and polymerized in a 60°C oven overnight. Images were acquired on a JOEL 1400 Plus transmission electron microscope equipped with a LaB6 source using a voltage of 120 kV, and the images were captured by an AMT BIOSPRINT 16M-ActiveVu mid mound CCD camera.

### Liver TAG measurement

Liver and serum samples were collected from WT and *Clstn3b-*LKO cohorts or WT and *Clstn3b* LTG cohorts maintained on HFD/WD. About 60 mg of liver was removed from each mouse for measuring TAG. TAG levels were measured with the Triglyceride Colorimetric Assay Kit (Cayman, 10010303) or Glycerol Assay Kit (Sigma, MAK117) following the manufacturer’s instructions.

### Proteomics

LD was isolated from mouse liver without protease digestion. The suspension was mixed with 10x volume of acetone and incubated at -20℃ overnight for delipidation and protein precipitation. The mixture was centrifuged at 12,000 g for 5 min. The pellets were washed with acetone and dried by heating at 60℃ for at least 15 min. Pellets were then washed with 20% TCA to remove excess sucrose, washed with acetone, and dried. The pellets were dissolved in 50 mM triethylammonium bicarbonate (TEAB, pH=8), 5% SDS at 60℃ with shaking for 30 min. Protein concentration was determined with a BCA method.

Samples were then reduced by adding dithiothreitol (DTT) to a final concentration of 10 mM and samples were incubated at 56°C for 30 min. After cooling, iodoacetamide was added to a final concentration of 20 mM and samples were alkylated for 30 min at room temperature in the dark. Following centrifugation for 2 min at 13.2 krpm, the supernatants were removed and digested overnight with trypsin at 37°C using an S-Trap (Protifi). Following digestion, the peptide eluate was dried and reconstituted in 100 mM TEAB buffer.

Samples were injected onto an Orbitrap Fusion Lumos mass spectrometer coupled to an Ultimate 3000 RSLC-Nano liquid chromatography system (Thermo). Samples were injected onto a 75 um i.d., 75-cm long EasySpray column (Thermo) and eluted with a gradient from 0-28% buffer B over 180 min. Buffer A contained 2% (v/v) ACN and 0.1% formic acid in water, and buffer B contained 80% (v/v) ACN, 10% (v/v) trifluoroethanol, and 0.1% formic acid in water. The mass spectrometer operated in positive ion mode with a source voltage of 1.8-2.0 kV and an ion transfer tube temperature of 300°C. MS scans were acquired at 120,000 resolution in the Orbitrap and top speed mode was used for SPS-MS3 analysis with a cycle time of 2.5 s. MS2 was performed with CID with a collision energy of 35%. The top 10 fragments were selected for MS3 fragmentation using HCD, with a collision energy of 55%. Dynamic exclusion was set for 25 s after an ion was selected for fragmentation.

Raw MS data files were analyzed using Proteome Discoverer v2.4 SP1 (Thermo), with peptide identification performed using a trypsin digest search with Sequest HT (cleavage after Lys and Arg except when followed by Pro). The mouse reviewed protein database from UniProt (downloaded Jan. 28, 2022, 17,062 entries) was used. Fragment and precursor tolerances of 10 ppm and 0.6 Da were specified, and three missed cleavages were allowed. A minimum peptide length of 6 residues was required. Carbamidomethylation of Cys and TMT10plex labelling of N-termini and Lys sidechains were set as a fixed modification, with oxidation of Met set as a variable modification. The false-discovery rate (FDR) cutoff was 1% for all peptides. At least two unique peptides were required for protein identification.

### Liver LD isolation and digestion

The liver tissue was dissected, minced into tiny pieces with a spring scissor and transferred to a motorized homogenizer in HES buffer (20mM HEPES + 1mM EDTA + 250mM Sucrose). The homogenate was filtered with double-layer gauze and centrifuged at 2000 g for 5 min. The infranatant was removed with a syringe and the buoyant LD fraction was transferred with a wide-opening tip into 5 ml tubes and washed with HES buffer 2 times. The LD was then transferred to Ultra-Clear ultracentrifuge tubes (Beckman-Coulter), adjusted to a final concentration of 20% sucrose, and overlaid by 5% sucrose/HE and HE (20mM HEPES + 1mM EDTA). The gradient was centrifuged at 16,000 g for 10 min at 4℃. The buoyant LD was then collected.

To remove the LD-bound organelles, the LD fraction was digested with 1 mg/mL proteinase K for 5-15 min at 37℃ and 1mM PMSF was added to inactivate proteinase K. The mixture was centrifuged through the a before mentioned sucrose gradient at 210,000g for 1 hr. The buoyant LD was then collected.

### Phospholipids quantitation

LD was isolated from mouse liver and digested with proteinase K following the procedure described above. The collected LD fractions (100-200 μL suspension in HES buffer) were transferred into round bottom glass tubes and extracted with a mixture of 1 mL hexane, 1 mL methyl acetate, 0.75 mL acetonitrile and 1 mL water as previously described. The extracts were vortexed for 5 s and centrifuge at 2,671 g for 5 min to partition into 3 phases. The upper and middle phases were collected into separate glass tubes and dried under N_2_. Phospholipids in the middle phase were measured with the Phosphatidylcholine Assay Kit (Colorimetric/Fluorometric) (Abcam, ab83377) and Phosphatidylethanolamine Assay Kit (Fluorometric) (Sigma, MAK361) following the manufacturer’s instructions. TG in the upper phase was dissolved in 350 μL ethanolic KOH (2 part EtOH and 1 part 30% KOH) and incubated overnight at 55°C for complete hydrolysis. The volume was then brought to 1200 μL with H2O: EtOH (1:1) and vortexed to mix. Two hundred μL was transferred to a new tube, mixed with 215 μL 1M MgCl_2_ and vortexed. The mixture was incubated on ice for 10 min and centrifuged at 13,000g for 5 min. Glycerol content in the supernatant was determined with the Free Glycerol Reagent (Sigma, F6428) following the manufacturer’s instructions.

To calculate LD surface phospholipids density, we normalized phospholipids abundance as measured by the fluorometric kit or phospholipidomics to LD surface area of each sample. The detailed normalization procedure is described in a previous publication (Zhang et al, PMID: 38293096). Briefly, we divided total TG content of each sample by the mean LD volume to derive total LD number, which was then multiplied by the mean LD surface area. Mean LD volume and surface area were determined by Imaris analysis of LD images harvested from the same batch of sample.

### Primary hepatocyte isolation and culture

Primary hepatocytes were isolated from mouse livers using a two-step collagenase perfusion method. Briefly, mice were anesthetized with isoflurane, and the abdominal area was sterilized with 70% ethanol. A midline incision was made to expose the portal vein, and a catheter was inserted into the inferior vena cava (IVC) and secured by threading a needle through the catheter’s butterfly wing. Perfusion was initiated with Liver Perfusion Medium (Life Technologies, #17701038) at a flow rate of 3 mL/min, using approximately 30–50 mL of medium. The portal vein was immediately severed to allow for adequate outflow. Following perfusion with Liver Perfusion Medium, the liver was perfused with Liver Digestion Medium (Life Technologies, #17703034) at the same flow rate, using an additional 30–50 mL. After digestion, the liver was carefully removed and placed in 10 mL of Liver Digestion Medium in a 10 cm culture dish. The liver capsule was gently peeled away, and the tissue was gently agitated to release hepatocytes. The cell suspension was then filtered through a 70 μm nylon mesh into a 50 mL tube prefilled with 20 mL of Hepatocyte Washing Medium (Life Technologies #17704024) on ice to halt enzymatic activity. Cells were pelleted by centrifugation at 50 g for 5 minutes at 4°C, and the supernatant was discarded. The cell pellet was resuspended in 20 mL of Washing Medium, and the wash step was repeated twice to ensure cell purity. Cell viability and yield were assessed by Trypan blue exclusion. Cells were then centrifuged once more at 50 g for 5 minutes and resuspended in the desired volume of washing medium or buffer for further experiments.

### Hepatocyte respiration

Hepatocyte oxygen consumption data were collected by high-resolution respirometry with an Oroboros Oxygraph-2k (Oroboros Instruments, Innsbruck, Austria) in a standard configuration, with 2 mL volume of the two chambers, at 37 °C, and 350 rpm stirrer speed. Specifically, freshly Isolated primary hepatocytes were suspended at a density of 1-2 × 10^6^ cells/mL in a respiration buffer consisting of 120 mM sodium chloride, 5 mM potassium chloride, 1 mM magnesium chloride, 1.3 mM calcium chloride, 0.4 mM dibasic potassium phosphate, 20 mM HEPES, 10 mM sodium bicarbonate, 10 mM glucose and 4% (w/v) BSA (fatty acid free) at pH 7.3, and same number of cells were added to the respirometry chamber for measurement. The software DatLab7 was used for data acquisition (2s time intervals) and analysis. All measurements were performed within 4 hours of cell isolation.

### Mitochondria isolation

All mitochondria isolation and fractionation were carried out on ice. Liver tissues were minced in ice-cold HES buffer (20 mM HEPES, 1mM EDTA, 250 mM sucrose, pH 7.2) supplemented with 1% fatty acid-free BSA (Millipore, Code82-002-4) and 1x EDTA-free Protease Inhibitor Cocktail (Thermo Scientific, 1861279), and gently homogenized with a Teflon pestle. Homogenates were passed through two layers of cheesecloth and centrifuged at 12,000 x g for 10 mins at 4°C. The fat layer was removed and discarded, and the resulting pellets were resuspended in ice-cold HES buffer. Samples were then centrifuged at 800 x g for 10 min at 4°C. The supernatants were then transferred to fresh tubes and centrifuged again at 1,300 x g for 10 min at 4°C. To achieve the mitochondrial fraction (pellet), the supernatants were again transferred to new tubes and centrifuged at 7000 x g for 10 min at 4°C. The resulting crude mitochondrial pellets were washed three times with 0.15 M KCl to remove catalase and then spun a final time in ice-cold HES buffer. The final mitochondrial pellets were resuspended in the relative buffer for experimental use.

### Anion-Exchange Chromatography for ATP and ADP measurement in hepatocytes

Primary hepatocytes cultured in 12 well plates were extracted with 500ul solvent containing Methanol:Acetonitrile:Water 2:2:1 (v/v/v). The extract was transferred to a 1.5ml Eppendorf tube and subjected to three freeze-thaw cycles between liquid nitrogen and 37°C water bath. After the third thaw, the samples were vortexed for 1 min and then centrifuged at a maximum speed for 15 minutes in a refrigerated centrifuge. The supernatant was transferred to a new tube and then dried in a SpeedVac to a volume just below 100ul. The sample was diluted to 500ul by adding 5 mM Tris-HCL 8.0, and resolved on Capto HiRes Q 5/50 0-40%B in 15ml (Buffer A: 5 mM Tris-HCl 8.0; Buffer B: 5mM Tris-HCl 8.0 and 1M NaCl). The ATP and ADP peaks were monitored and quantified based on UV absorption at 254 nm.

### Measurement of mitochondrial H_2_O_2_ production

Mitochondrial hydrogen peroxide (H_2_O_2_) production was measured fluorometrically in isolated liver mitochondria using the Amplex UltraRed/horseradish peroxidase (HRP) assay. Isolated mitochondria were first treated with 35 μM 1-chloro-2,4-dinitrobenzene (CDNB) for 5 min at room temperature, followed by three washes with ice-cold HES buffer to remove residual reagent. For the assay, a reaction mixture (200 μL per well) was prepared containing reaction buffer (125 mM KCl, 10 mM HEPES, pH 7.4, 2 mM MgCl₂, 2 mM KH_2_PO_4_, and 0.1% fatty acid-free BSA) supplemented with 10 μM Amplex UltraRed (Invitrogen), 3 U/mL HRP, and 20 U/mL Cu/Zn superoxide dismutase (SOD). The reaction mixture was equilibrated at 37 °C for 5 min before the addition of mitochondria. Mitochondrial suspensions were then added to achieve a final concentration of 50 μg protein per well, mixed gently, and baseline fluorescence was recorded for 5 min. Fluorescence measurements were performed using a Varioskan Lux plate reader (Thermo Fisher Scientific) at excitation/emission wavelengths of 565/600 nm. Mitochondrial H_2_O_2_ production was initiated by the addition of succinate (10 mM) directly to the wells (without removing the plate; add gently using an automated pipette, volume ≤ 10 µL). Fluorescence was recorded for an additional 5 min. H_2_O_2_ standards prepared in the same reaction buffer were included on each plate to allow quantification of H_2_O_2_ production rates from the linear increase in resorufin fluorescence.

### Measurement of Mitochondrial ROS

Mitochondrial reactive oxygen species (mitoROS) in primary hepatocytes were measured using a mitochondria-targeted fluorescent probe (MitoROS™ Red dye; Thermo Fisher Scientific; M36008). Cells were washed with pre-warmed PBS and incubated with MitoROS dye (500 nM) in serum-free medium for 30 min at 37°C in the dark. Wash cells gently 3 times with warm buffer (HBSS with Calcium and Magnesium or suitable buffer). After washing, fluorescence was immediately analyzed by fluorescence microscopy using appropriate excitation/emission settings (Ex/Em ≈ 540/590 nm). Mean fluorescence intensity was quantified by ImageJ/Fiji.

### Clear native PAGE and Coomassie staining for analysis of mitochondrial respiratory chain supercomplexes

Mitochondrial respiratory chain supercomplexes were analyzed by clear native polyacrylamide gel electrophoresis (CN-PAGE) followed by Coomassie staining. Briefly, Isolated mitochondria (50 μg) were solubilized in 20 μL sample buffer (5 μL of 4x Native Page Sample Buffer, 4 μL 10% digitonin, 11 μL ddH2O per sample) for 20 min on ice and then centrifuged at 20,000 x g for 30 mins at 4°C. 15 μL of the supernatant was collected and placed into a new tube, and mixed with 2 μL of G-250 sample buffer additive. Light blue cathode buffer (1 mL 20X Native Page running buffer, 10 mLs 20x cathode additive, 189 mLs ddH_2_O) was carefully added to the front of gel box (Invitrogen Mini Gel Tank A25977) and anode buffer (50 mLs 20x Native Page running buffer to 950 mL ddH_2_O) was carefully added to the back of the gel box making sure to not mix. The samples were then loaded onto a native PAGE 3-12% Bis-Tris Gel (BN1001BOX, Thermo Fisher Scientific), and electrophoresis was performed at 150 V for 30 min on ice. The light blue cathode buffer was carefully replaced with anode buffer and run at 250 V for 150 min on ice. Following electrophoresis, gels were fixed and stained with Coomassie Brilliant Blue R-250 to visualize native respiratory chain complexes and supercomplexes. Band patterns corresponding to individual complexes and higher-order supercomplex assemblies were documented by gel imaging and quantified by densitometry using ImageJ. Where indicated, gel bands were further processed for in-gel activity assays. All experiments were performed with at least three independent biological replicates.

### Complex I In-Gel Activity Assay

Mitochondrial complex I activity was assessed by an in-gel activity assay following Blue Native polyacrylamide gel electrophoresis (BN-PAGE). After electrophoresis, gels were equilibrated in assay buffer (25 mM Tris-HCl, pH 7.4) for 10 min at room temperature. Gels were then incubated at room temperature in freshly prepared complex I activity staining solution containing 2 mM Tris-HCl (pH 7.4), 0.1 mg/mL NADH, and 2.5 mg/mL nitrotetrazolium blue chloride (NTB) dissolved in distilled water. The reaction was allowed to proceed until distinct purple formazan bands corresponding to NADH dehydrogenase (complex I) activity became visible. The reaction was terminated by incubation in 10% acetic acid, followed by extensive washing with distilled water. Gels were imaged immediately, and complex I activity was quantified by densitometric analysis of band intensities using ImageJ/Fiji.

### Copper-free click chemistry phospholipid labeling and FRAP

U2OS was cultured for 12 h and then treated with OA (100 μM) or OA+LA(50 μM each)and azide choline (50 μM) for 12-24 h to allow LD formation. The cells were then washed with PBS twice and transfected with pCDNA3.1 (empty vector), CLSTN3B-mCherry, or RK-mCherry overnight or 12 hrs. The short duration was used to restrict CLSTN3B/RK expression and prevent complete ER wrapping around LDs. The cells were washed with PBS twice and incubated in serum-free DMEM with BDP FLDBCO (100nM) for 15min. The cells were then washed with fresh growth medium and allowed to sit for 60 min before the start of imaging. Regions of interest were bleached by 100% laser power (488 diode laser), followed by time-lapse imaging with a 23-second interval. The fluorescence intensity of BDP FL DBCO on the LD monolayer was quantified by ImageJ/Fiji.

### Alkynyl-linoleic acid/oleic acid labeling in LD-associated ER

HEK293 cells were transfected with Sec61β-GFP together with CLSTN3B-BFP or an empty vector control and labeled with alkynyl analogs of linoleic acid (LA-alk; Cayman Chemical, 10541) or oleic acid (OA-alk; Cayman Chemical, 9002078). Briefly, cells were incubated with LA-alk or OA-alk at a final concentration of 25 μM in the presence of 100 μM unlabeled oleic acid to induce LD biogenesis for 12 h. Following labeling, cells were washed with ice-cold phosphate-buffered saline (PBS), fixed with 4% paraformaldehyde, and permeabilized with 25 μM digitonin. Alkynyl fatty acids were visualized by copper(I)-catalyzed azide–alkyne cycloaddition (CuAAC) using azide-conjugated fluorophores. Briefly, cells were incubated for 1 h at room temperature in click reaction solution containing 50 μM Alexa Fluor™ 555 Azide (A20012, Thermo Fisher Scientific), a BTTAA(Click chemistry tools, 1236-100)–CuSO₄ complex (50 mM CuSO₄; BTTAA/CuSO₄, 6:1 molar ratio), and 2.5 mM sodium ascorbate. Neutral lipid droplets were stained with HCS LipidTOX™ Deep Red Neutral Lipid Stain (1:200; H34477, Thermo Fisher Scientific) for 15 min. Confocal images were acquired using a ZEISS LSM 900 microscope.

### Time-resolved analysis of mitochondrial enrichment of alkynyl-linoleic acid

HEK293 cells were transfected with mitochondria-targeting GFP together with CLSTN3B-BFP or an empty pcDNA3.1 vector. Twenty-four hours after transfection, cells were labeled with an alkynyl analog of linoleic acid (LA-alk; Cayman Chemical, 10541). Cells were incubated with LA-alk at a final concentration of 25 μM in the presence of 100 μM unlabeled oleic acid to induce lipid droplet biogenesis for 1, 3, 6, or 12 h. Following labeling, cells were visualized by click chemistry system using azide-conjugated fluorophore Alexa Fluor™ 555 Azide (A20012, Thermo Fisher Scientific), as described before. Confocal images were acquired using a ZEISS LSM 900 microscope. Mitochondrial enrichment of LA-alk was quantified by measuring the spatial overlap between LA-alk fluorescence and mito-GFP-labeled mitochondria with Imaris 11.0

### Image analysis

All images were taken on a ZEISS 900 confocal microscope with Airyscan. To analyze isolated lipid droplets, images were imported to Imaris 10.2.0 (Bitplane). Individual LDs were rendered by the “Surface” function and surface area and volume were measured. The rendering parameters were kept the same between treatments in each experiment. To analyze the extent of CLSTN3B wrapping of LDs, individual LDs and the surrounding CLSTN3B signals were rendered by the “Surface” function. The fraction of the volume bound by the LD surface overlapping with the volume bound by the CLSTN3B signal surface was calculated with the “Object-Object statistics” function. All plots were generated in R. Immunofluorescence staining for aSMA and F4/80 was performed on frozen liver sections, and fluorescence signal was analyzed using ImageJ (NIH). Images were captured under identical exposure settings using ZEISS 900 confocal microscope. Red fluorescence signals (representing aSMA or F4/80) were extracted by splitting color channels, and a threshold was applied to define positive signal areas. The percentage of aSMA or F4/80 area was calculated as the ratio of red-positive area to total tissue area. All signal analyses were performed in ImageJ.

To quantify the recovery of DBCO signal on lipid droplets (LDs), regions of interest (ROIs) with identical areas were first selected on LDs to measure DBCO fluorescence intensity prior to laser bleaching. The same ROIs were then applied to the identical LD positions at time 0 immediately after bleaching and at the final time point. For normalization, DBCO fluorescence intensity was measured at each corresponding time point in equal-area ROIs on adjacent ER regions that were not affected by bleaching. The recovery of DBCO signal on LDs was calculated as the ratio of normalized fluorescence intensity at the final time point to that measured at the same LD position before bleaching.

To assess enrichment of LA-Alk or OA-alk on lipid droplet–associated ER, equal-area regions of interest (ROIs) were defined on ER membranes adjacent to lipid droplets and on non–lipid droplet–associated ER. Alk fluorescence intensity was quantified using ImageJ, and enrichment was calculated as the ratio between lipid droplet-associated ER and non-lipid droplet-associated ER signal intensity

To analyze mitochondria-associated LA-alkyne signal level, images were imported into Imaris 11.0 and mitochondria were segmented by the “Surface” function using the mito-GFP channel. A mask was then created to set all pixels outside the surface to zero. LA-alkyne signal was then rendered by the “Surface” function using the Alexa Fluor™ 555 Azide channel and total signal intensity enclosed by the surface was computed.

### Real-time qPCR analysis

The following primers were used for qPCR analysis of gene expression. *Clstn3b*-fwd, CTCCGCAGGAACAGCAGCCC, rev, AGGATAACCATAAGCACCAG; *tnf*-fwd, GGTGCCTATGTCTCAGCCTCTT, rev, GCCATAGAACTGATGAGAGGGAG; *ccl2*-fwd, GCTACAAGAGGATCACCAGCAG, rev, GTCTGGACCCATTCCTTCTTGG; *adgre1*-fwd, CGTGTTGTTGGTGGCACTGTGA, rev, CCACATCAGTGTTCCAGGAGAC; *lipe*-fwd, TTGGGGAGCTCCAGTCGGA, rev, TCGTGCGTAAATCCATGCTGT; *fabp4*-fwd, ACACCGAGATTTCCTTCAAACTG, rev, CCATCTAGGGTTATGATGCTCTTCA; *L19*-fwd, GGTCTGGTTGGATCCCAATG, rev, CCCATCCTTGATCAGCTTCCT; *lpl*-fwd, GGGAGTTTGGCTCCAGAGTTT, rev, TGTGTCTTCAGGGGTCCTTAG; *cfd*-fwd, CATGCTCGGCCCTACATGG, rev, CACAGAGTCGTCATCCGTCAC; pparg-fwd, GTACTGTCGGTTTCAGAAGTGCC, rev, ATCTCCGCCAACAGCTTCTCCT; ppara-fwd, AGAGCCCCATCTGTCCTCTC, rev ACTGGTAGTCTGCAAAACCAAA; ppargc1a-fwd, GAATCAAGCCACTACAGACACCG, rev, CATCCCTCTTGAGCCTTTCGTG; cpt2-fwd, CAGCACAGCATCGTACCCA, rev,TCCCAATGCCGTTCTCAAAAT cpt1a-fwd, CTCCGCCTGAGCCATGAAG, rev,CACCAGTGATGATGCCATTCT; acads-fwd, TGGCGACGGTTACACACTG, rev,GTAGGCCAGGTAATCCAAGCC; acadl-fwd, TCTTTTCCTCGGAGCATGACA, rev, GACCTCTCTACTCACTTCTCCAG; acadm-fwd, AGGGTTTAGTTTTGAGTTGACGG, rev, CCCCGCTTTTGTCATATTCCG; acaa2-fwd, GAATCAAGCCACTACAGACACCG,rev, CATCCCTCTTGAGCCTTTCGTG; etfa-fwd, GTCTTGGAGGTGAAGTGTCCTG, rev, GCATCATGCTGAGCCACCAGAA ; etfdh-fwd, ATGGAGGCTCTTTCCTTTACCAC rev, GCTTACACCTCTGGAACTCTCG ; echs1-fwd, CTCAACCAAGCACTGGAGACCT rev, GCTGGAGTAACAGTCCTGAAATG ; hadhb-fwd, CACTGCGTTCTCATAGTCTGGC rev, GCCATTTGCTCCAGTGAGGAAG; hadha-fwd, GTTTGAGGACCTCGGTGTAAAGC rev, GAGAGCAGATGTGTTGCTGGCA; tgfb1-rwd, TGATACGCCTGAGTGGCTGTCT, rev, CACAAGAGCAGTGAGCGCTGAA; timp1-fwd, TCTTGGTTCCCTGGCGTACTCT rev, GTGAGTGTCACTCTCCAGTTTGC; acat2-fwd, GAGATTGTGCCAGTGCTGGTGT, rev, GTGACAGTTCCTGTCCCATCAG ; cola1-fwd, CCTCAGGGTATTGCTGGACAAC, rev, CAGAAGGACCTTGTTTGCCAGG ; mmp2-fwd, CAAGGATGGACTCCTGGCACAT, rev, TACTCGCCATCAGCGTTCCCAT; mmp9-fwd, GCTGACTACGATAAGGACGGCA, rev, TAGTGGTGCAGGCAGAGTAGGA; desmin-fwd, GCGGCTAAGAACATCTCTGAGG, rev, ATCTCGCAGGTGTAGGACTGGA; pdgfrb-fwd, GTGGTCCTTACCGTCATCTCTC, rev, GTGGAGTCGTAAGGCAACTGCA; tnfa-fwd, GGTGCCTATGTCTCAGCCTCTT, rev, GCCATAGAACTGATGAGAGGGAG; bax-fwd, AGGATGCGTCCACCAAGAAGCT, rev, TCCGTGTCCACGTCAGCAATCA. actin-fwd, CATTGCTGACAGGATGCAGAAGG, rev, TGCTGGAAGGTGGACAGTGAGG.

### ChIP-qPCR

The liver tissue was dissected, minced into tiny pieces with a spring scissor and transferred to a motorized homogenizer in HES buffer (20mM HEPES + 1mM EDTA + 250mM Sucrose). Add 32% PFA into homogenizing mixture to final 1% concentration and crosslink by gently rocking at room temperature for 10 minutes, followed by quenching with 0.125 M glycine. Nuclei were collected by centrifugation and lysed with nuclei lysis buffer (1% SDS, 50 mM Tris-HCl, 10 mM EDTA at pH 8.0) and sonicated to shear DNA to ∼100–500 bp fragments. The lysate was diluted with a dilution buffer (0.11% sodium deoxycholate, 1.1% Triton X-100, 50 mM Tris-HCl, 167 mM NaCl at pH 8.0) and kept some lysate to yield input DNA. Incubate the diluted lysate with anti-PPARγ antibody (Proteintech, Cat#6643-1-AP) overnight at 4°C. The protein–DNA complex was isolated with protein G Dynabeads, washed, and eluted off the beads by elution buffer (0.5% SDS, 10 mM Tris-HCl pH8, 5 mM EDTA at pH 8.0). The eluent was treated with RNase A and proteinase K before reversing crosslinking. DNA was then harvested by ethanol precipitation and analyzed by qPCR using primers targeting PPARγ-binding regions and non-binding regions in selected gene promoters. Input was used as control.

### Metabolic cage

Energy expenditure and metabolic parameters were assessed using a comprehensive lab animal monitoring system (CLAMS; Columbus Instruments) and operated by the UT Southwestern Metabolic Phenotyping Core. Mice were singly housed in metabolic cages for continuous monitoring of oxygen consumption (VO₂), carbon dioxide production (VCO₂), respiratory exchange ratio (RER), locomotor activity, food intake, and water consumption. Animals were acclimated in the chambers for 24 hours prior to data collection. Measurements were recorded over a 45-hour period under a 12-hour light/dark cycle at 30°C with ad libitum access to food and water. Data were analyzed using the manufacturer’s software and normalized to body weight.

### Temperature measurement

Core body temperature of mice was measured using a rectal thermometer (BAT-12, PHYSITEMP) with a lubricated probe inserted ∼2 cm into the rectum. Mice were housed in climate Chamber at 30 . Mice were gently restrained by hand, and temperature was recorded once a stable reading was obtained, typically within 10-15 seconds. All measurements were performed at the same time of day at midnight. To assess surface temperature over specific anatomical regions by infrared thermograph, mice were anesthetized briefly, and hair was removed from the skin overlying the liver (right upper abdominal area) and interscapular brown adipose tissue (BAT) using hair Clipper and the treat with commercial depilatory cream. After full recovery from anesthesia, mice were placed individually in a temperature-controlled environment, and thermal images were captured using an infrared camera (TC004, TOPDON TECH). Surface temperatures over the liver and BAT regions were quantified using ImageJ to measure the signal intensity and compared with the temperature bar to calculate the corresponding temperature.

### FAO blue staining

Fatty acid oxidation (FAO) activity was assessed using the FAO Blue™ kit (FNK-FDV-0033, Diagnocine) according to the manufacturer’s instructions. Briefly, cells were incubated with the 20uM FAO Blue working solution at 37°C for 30 minutes in the dark. After staining, cells were washed with the PBS and changed to fresh medium without serum and subjected to live-cell imaging using a fluorescence microscope (excitation/emission: 405/450 nm).

### TMRE staining

Mitochondrial membrane potential staining was performed using TMRE (I34361, Thermo Fisher) following the product instruction. Briefly, cells were incubated with 100 nM TMRE diluted in culture medium at 37°C for 30 minutes. After incubation, cells were immediately subjected to live-cell imaging using a fluorescence microscope (excitation/emission: 548/574 nm) without washing.

### LD size distribution plotting

To visualize the relative volume distribution of LDs, a volume-weighted density graph was plotted in which the x value is LD volume (log transformed) and the y value is the density weighted by the relative percentage of total LD volume. To characterize the volume-weighted volume distribution, a probability function was first generated from the density distribution by the “approxfun” function in R. The median value of the distribution was then calculated by random simulation (“simulate” function, 1e6 times) of the probability function in R.

### Lipidomics analysis

The solvents used were either HPLC or LC/MS grade and purchased from Sigma-Aldrich (St Louis, MO, USA). Splash Lipidomix stadndards were purchased from Avanti (Alabaster, AL, USA). All lipid extractions were performed in 16×100mm glass tubes with PTFE-lined caps (Fisher Scientific, Pittsburg, PA, USA). Glass Pasteur pipettes, eVol automated analytical syringes (Trajan, Australia) and solvent-resistant plasticware pipette tips (Mettler-Toledo, Columbus, OH, USA) were used to minimize leaching of polymers and plasticizers. The lipid liquid-liquid extraction was performed using a modified methyl-tert-butyl ether (mTBE) method (Matyash et al., PMID: 18281723). Briefly, samples were transferred to a glass tube where 1mL of water, 1mL of methanol and 2 mL of mTBE were added. The mixture was vortexed, then centrifuged at 2671×g for 5 min, the organic phase was collected, spiked with 20µL of a 1:5 diluted Splash Lipidomix standard solution and dried under N2 air flow. The samples were resuspended in hexane for lipidomics analysis. The lipid profile was obtained by LC-MS/MS using a SCIEX QTRAP 6500+ (SCIEX, Framingham, MA) equipped with a Shimadzu LC-30AD (Shimadzu, Columbia, MD) high-performance liquid chromatography (HPLC) system and a 150×2.1 mm, 5 µm Supelco Ascentis silica column (Supelco, Bellefonte, PA). Samples were injected at a flow rate of 0.3 ml/min at 2.5% solvent B (methyl tert-butyl ether) and 97.5% Solvent A (hexane). Solvent B was increased to 5% over 3 min and then to 60% over 6 min. Solvent B was decreased to 0% during 30 sec while Solvent C (90:10 (v/v) isopropanol-water) was set at 20% and increased to 40% during the following 11 min. Solvent C is increased to 44% over 6 min and then to 60% over 50 sec. The system was held at 60% solvent C for 1 min prior to re-equilibration at 2.5% of solvent B for 5 min at a 1.2 mL/min flow rate. Solvent D [95:5 (v/v) acetonitrile-water with 10 mM Ammonium acetate] was infused post-column at 0.03 ml/min. Column oven temperature was 25°C. Data was acquired in positive and negative ionization mode using multiple reaction monitoring (MRM). The LC-MS/MS data was analyzed using MultiQuant software (SCIEX). All samples were normalized to protein content and identified lipid species were normalized to its corresponding internal standard.

### Liposome preparation

Large unilamellar vesicles (LUV) were prepared as described before (Yang et al., Biophys. J., 2010). Lipid mixtures dissolved in a benzene/methanol mixture (9:1 ratio) were frozen in liquid nitrogen and freeze-dried overnight in a CentriVap vacuum concentrator (Labconco, USA). The dried lipid was then resuspended in a buffer (100 mM NaCl, 10 mM Hepes, 5 mM EGTA, pH 7.0) total lipid concentration of 0.5 mM. The suspension was passed ten times through the double-stacked nucleopore polycarbonate track etch membrane filters with a nominal pore size of 0.1 µm using a Lipex liposome extruder. Liposomes were used within 1 day.

### Lipid mixing experiments

Lipid mixing was measured by the release of fluorescence resonance energy transfer between TopFluor PE (0.5 mol %) and rhodamine PE (0.25 mol %). Unlabeled liposomes of various lipid compositions and liposomes labeled with a self-quenching concentration were added to 2 mL of buffer in quartz cuvette in a ratio of 10:1 to a total lipid concentration of 10 μM. Different concentrations of peptides were then added to induce lipid mixing between liposomes. Lipid mixing between labeled and unlabeled liposomes results in a dilution of fluorescent probes and an increase in dye fluorescence due to a relief of self-quenching. Fluorescence changes induced by peptides were recorded under constant stirring using a PC1 photon-counting spectrometer (ISS, USA) with λex = 480 nm and λem = 505 nm. At the end of each recording, complete dequenching of the dye was induced by adding Triton X-100 (0.1% v/v final concentration). All experiments were performed at 37 °C. The degree of dye quenching was calculated as Q(t) = 100 × (F(t) - F0)/(Ftriton - F0), where F(t), F0, and Ftriton are fluorescence at time t, before the addition of peptide and after addition of Triton X-100, respectively. The initial rate (v0) and final extent (A0) of lipid mixing were estimated from the fit of the Q(t) curve by the expression Q(t) = A0 - A0* exp(-(t-t0)/τ) with v0 = A0/τ.

### Lipid strip binding assay

The interaction between CLSTN3B and phosphatidic acid (PA) was examined using a lipid overlay assay with Cayman LipiDOT Strips™ membranes (38924, Cayman Chemical). Membranes were first blocked in Tris-buffered saline (TBS) containing 3% fatty acid–free bovine serum albumin (BSA) for 12 h at 4 °C to minimize nonspecific binding. Purified CLSTN3B protein (4 μg/mL), isolated from HEK293 cells expressing CLSTN3B as indicated, was diluted in blocking buffer and incubated with the lipid strips for 1 h at room temperature with gentle agitation. Following incubation, membranes were washed 5-6 times with wash buffer (≥10 mL per wash, 10 min each) at room temperature with gentle agitation. Membranes were then incubated with the appropriate primary antibody diluted in blocking buffer for 1 h at room temperature, followed by 5-6 washes as described above. Subsequently, membranes were incubated with the appropriate secondary antibody diluted in blocking buffer for 1 h at room temperature, washed 5-6 times, and protein-lipid interactions were detected by chemiluminescence.

### LNP preparation

Lipid nanoparticles were prepared by the ethanol dilution method as described previously (Vaidya et al., PMID: 38973655; Wang et al., PMID: 36316378; Cheng et al., PMID: 32251383). Lipids (5A2SC8/DOPE/Cholesterol/DMG-PEG/DODAP) were dissolved in ethanol and siRNA [Scrambled (Control) and Clstn3b-targeting (siClstn3b)] was diluted in 10 mM citrate buffer, pH 4.0. The internal molar fraction of the five lipid components was maintained at 15/15/30/3/15.75 (mol/mol). As a representative example, LNPs encapsulating 10 μg of siRNA were formulated with 6.62 mM 5A2-SC8, 6.62 mM DOPE, 13.23 mM Cholesterol, 1.32 mM DMG-PEG, and 6.95 mM DODAP resulting in a final molar ratio of 19.05/ 19.05/ 38.10/ 3.8/ 20 (mol/mol). The total lipid : siRNA ratio (w/w) was 40:1. Lipid (ethanol phase) and siRNA (aqueous phase) was rapidly mixed with a pipette in 3:1 ratio (v/v) to obtain LNPs. LNPs were dialyzed against 1× PBS for at least 2 h (Pur-A-Lyzer Midi 3500 Dialysis Kits, MWCO 100 kDa, Sigma) and diluted suitably with PBS prior to use. LNPs were ∼ 170 nm in size with a low PDI (0.17) which was measured with a Zetasizer Nano ZS dynamic light scattering (DLS) instrument (Malvern Instruments; He−Ne laser, wavelength = 632 nm). For the in vivo knockdown experiments related to reversal of MASLD pathology, mice were injected with two doses of 0.5 mg/kg separated by 48 h, and liver was harvested after two weeks for further analysis.

### MASLD patient cohorts

Associations of hepatic CLSTN3B expression with clinical phenotypes/outcomes and molecular pathways were assessed in four previously published cohorts of MASLD patients with hepatic transcriptome data (Govaere et al, PMID: 33268509; Fujiwara et al, PMID: 35731891; Hong et al, PMID: 31467298; Pantano et al, PMID: 34508113). The first cohort includes 206 European Caucasian patients with MASLD (histological fibrosis stage, F0-4) and 10 healthy obese controls (NCBI Gene Expression Omnibus, accession number GSE135251). The second cohort includes 106 Japanese patients with MASLD (F0-3 fibrosis), among which 77 patients underwent paired liver biopsies with a median interval of 2.3 years and transcriptome profiling (GSE193084). The third cohort include 72 MASLD patients (F0-4 fibrosis) and 6 healthy controls in the U.S. (GSE130970). The fourth cohort includes 112 MASLD patients (F0-4 fibrosis) and 31 healthy controls in the U.S. (GSE162694).

### Quantitation of CLSTN3B gene expression level

Raw sequencing reads in FASTQ format were preprocessed using Cutadapt (v2.5) (Martin, M. Cutadapt removes adapter sequences from high-throughput sequencing reads. EMBnet.journal 17, 10–12 (2011)) to trim low-quality bases and adapter sequences (parameters “-q 20 --max-n 3 -a Ill_Univ_Adapt”). The trimmed reads were then aligned to the human reference genome (GRCh38.p14) using STAR aligner (v2.7.3a) (Dobin et al, PMID: 23104886) with parameters, “--chimSegmentMin 15 --chimJunctionOverhangMin 15”. Gene-level quantification was performed using featureCounts (v1.6.3) (Liao et al, PMID: 24227677), assigning aligned reads to annotated gene features. Expression levels of CLSTN3B and CLSTN3 genes were quantified using exons unique to each transcript based on ENST00000535313.2 from the GENCODE release 45 annotation. Raw read counts were normalized using the relative log expression (RLE) method implemented in the DESeq2 R package (v1.42.0) (Love et al, PMID: 25516281). Log-transformed (base 2) RLE data were used for downstream analyses. Association of CLSTN3B and CLSTN3 expression and molecular pathway dysregulation was assessed using gene set enrichment analysis (www.gsea-msigdb.org/gsea). Bioinformatic data analysis was performed using R and python languages on the BioHPC supercomputing facility at University of Texas Southwestern Medical Center (portal.biohpc.swmed.edu).

## Acknowledgement

We thank Dr. Leonid Chernomordik for discussion on the liposome fusion and phospholipid mixing assay; Dr. Xun Wang at Children’s Research Institute at UTSW for the guidance on primary hepatocyte isolation from mice on HFD; Dr. Andrew Lemoff and the Proteomics core at UTSW for the proteomics analysis; the Molecular Pathology core at UTSW for the histology analysis; the EM core at UTSW (supported by NIH grant 1S10OD021685-01A1) for project discussion and EM sample preparation. The automated bioinformatics pipeline was developed by J.C. of the Data Science Shared Resource (DSSR) at UTSW Simmons Comprehensive Cancer Center (SCCC). Bioinformatics analyses were conducted on BioHPC, High-Performance Computing (HPC) resources embedded in Lyda Hill Department of Bioinformatics at UTSW. X.Z. is a Rita C. and William P. Clements, Jr. Scholar in Biomedical Research. This study was supported by the Endowed Scholars in Medical Science Program at UTSW, Cancer Prevention and Research Institute of Texas grant RR200084, NIH R01DK135556, and American Heart Association Award 23CDA1050474 to X.Z. Y.H. is supported by U.S. National Institutes of Health (CA233794, CA255621, CA282178, CA288375, CA283935); European Commission (ERC-AdG-2020-101021417); Cancer Prevention and Research Institute of Texas (RR180016, RP200554). H.S. is supported by Uehara Memorial Foundation. J.C. and J.L. are partly supported by the UTSW Simmons Comprehensive Cancer Center (SCCC) grant P30CA142543. A.S. and K.M. were supported in part by the Intramural Research Program of the National Institutes of Health (NIH). The contributions of the NIH author(s) are considered Works of the U.S. Government. The findings and conclusions presented in this paper are those of the author(s) and do not necessarily reflect the views of the NIH or the U.S. Department of Health and Human Services. D.J.S. acknowledge financial support from the National Institutes of Health (NIH) National Institute of Biomedical Imaging and Bioengineering (NIBIB) (R01 5R01EB025192-06).

## Author contributions

C.Z. and X.Z conceived the project. C.Z. and X.Z. designed the experiments. C.Z., D.Y., M.L., and X.Z. performed the bulk of the experiments. G.D.V. and J.G.M. conducted lipidomics analyses. H.S., J.C., J.L., and Y.H. analyzed the association between human *CLSTN3B*/*CLSTN3* expression and MASLD parameters. J.W., Q.Z., and H.Z. provided the MOSAICS system for AAV-mediated gene deletion in hepatocytes, prepared AAV for exogenous gene expression in the liver, and provided guidance on establishing primary hepatocyte cultures. K.M. and A.S. performed the FRET-based lipid mixing assay. A.V. and D.J.S. provided the LNP/siRNA system for acute hepatic *Clstn3b* knockdown and prepared the LNP/siRNA mixture. M.Y. performed part of the image and statistical analyses. J.Z. performed liver cryosections and immunostaining for fibrosis analysis and helped with LNP injection. X.Z. and C.Z. wrote the manuscript. All authors participated in reviewing and discussing the manuscript.

## Competing interests

Y.H. owns stock in Alentis Therapeutics and Espervita Therapeutics, and advises on Helio Genomics, Espertiva Therapeutics, Roche Diagnostics, Elevar Therapeutics.

---

**Figure S1.**
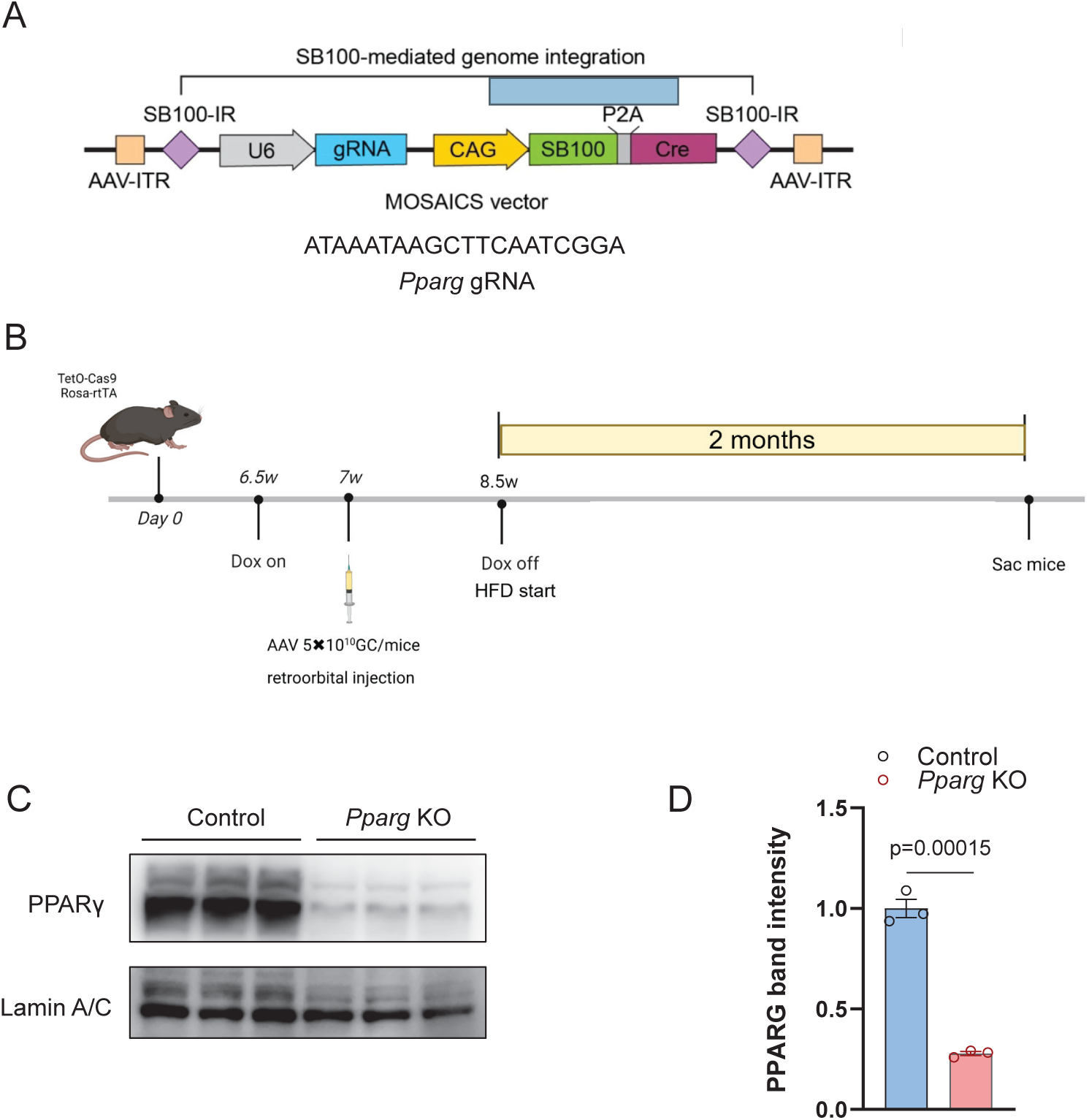
**A**, Schematic diagram of the MOSAICS vector containing *Pparg* gRNA for packaging adenovirus. **B**, *Pparg* KO mouse construction and HFD induction timeline. **C**-**D**, western blot analysis (**C**) and quantitation (**D**) of PPARY expression in nuclear extracts isolated from liver of control or *Pparg* KO mice on HFD (n=3). Data are mean ± s.e.m. Statistical significance was calculated by unpaired Student’s two-sided t-test in **D**.

**Figure S2.**
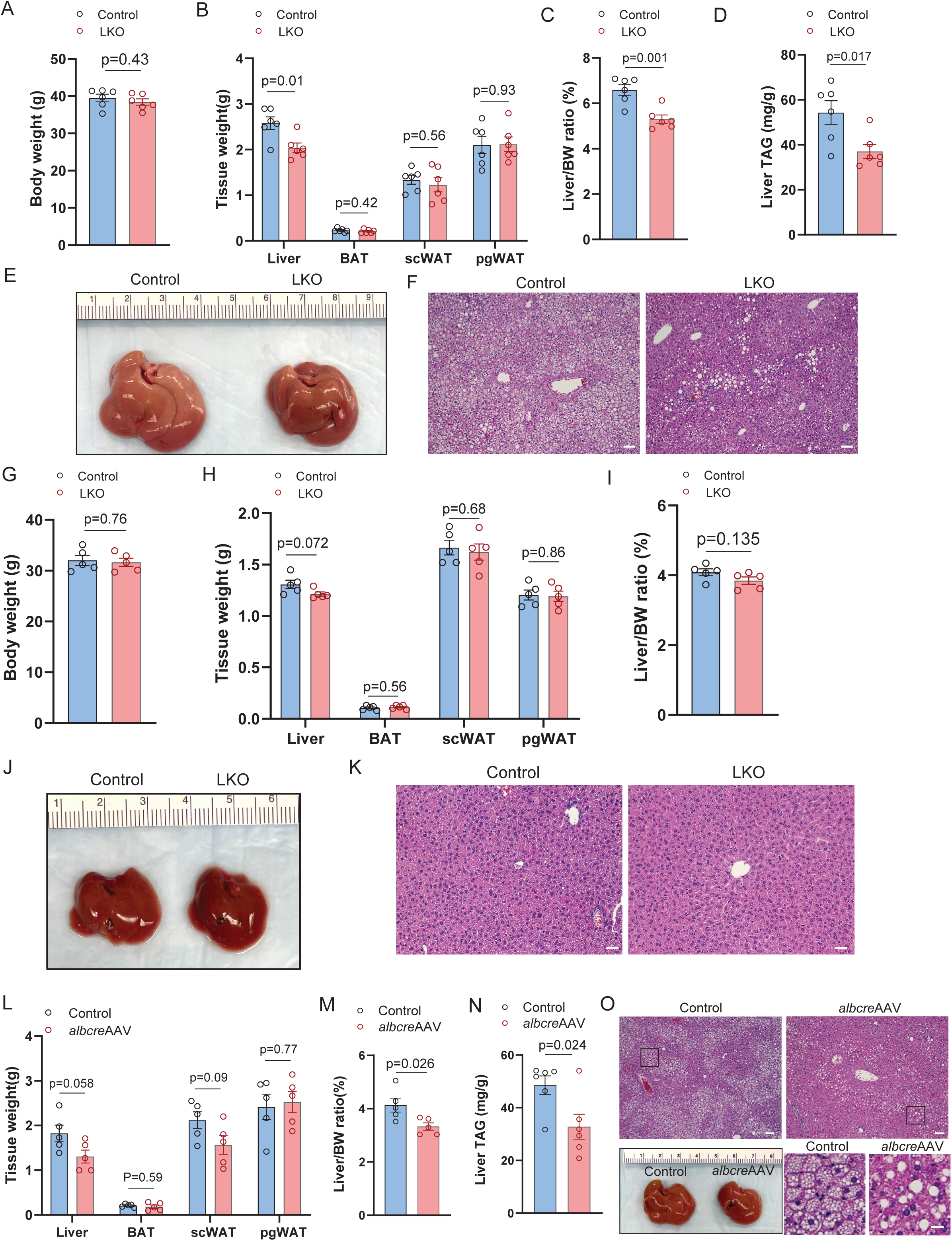
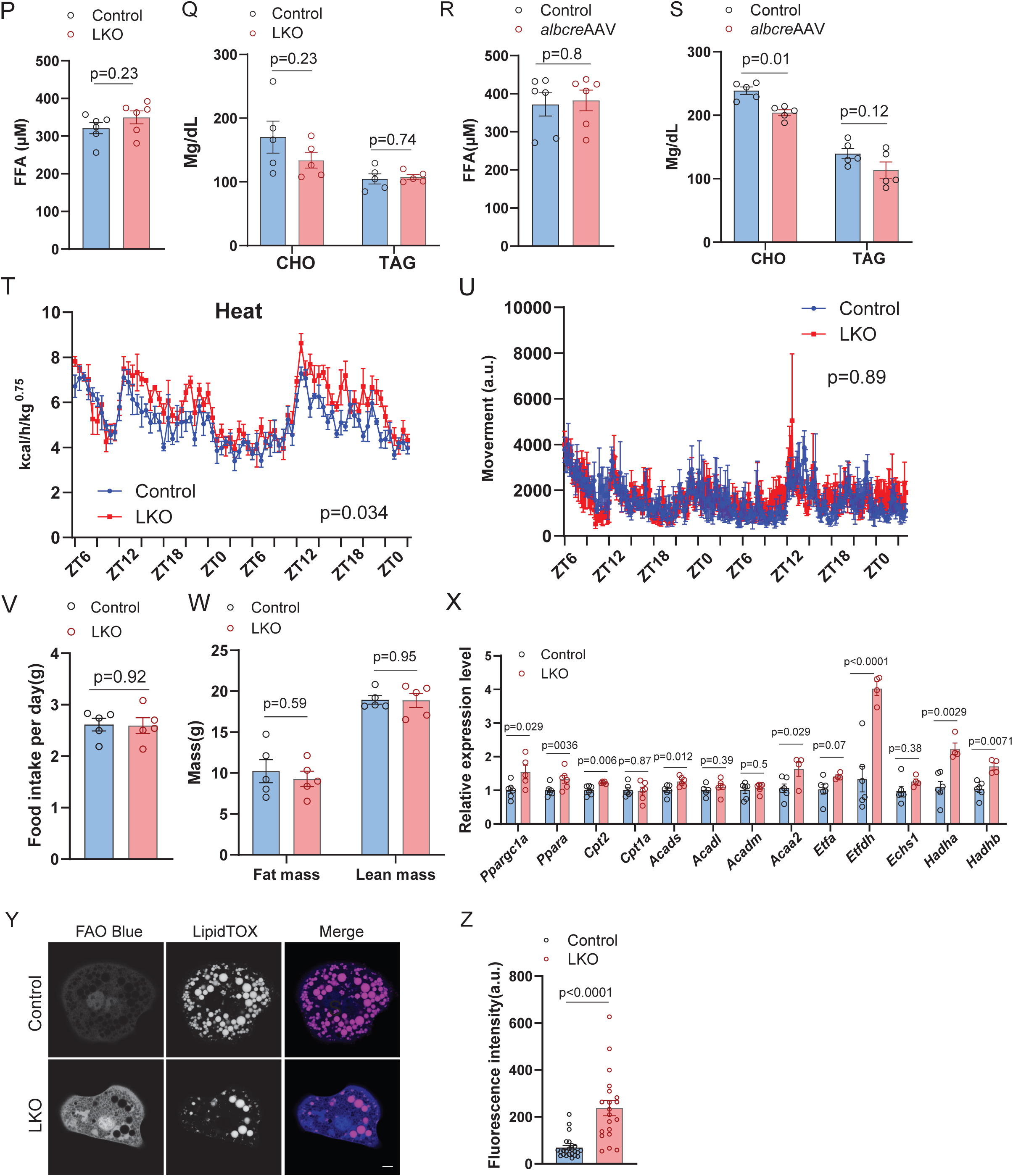
**A**-**F**, Body weight (**A**), tissue weights (**B**), liver/body weight ratio (**C**), liver TAG content (**D**), liver gross appearance (**E**) and histology (**F**) from WT control and LKO mice on WD for 10 weeks (n=5). **G**-**K**, Body weight (**G**), tissue weights (**H**), liver/body weight ratio (**I**), liver gross appearance (**J**) and histology (**K**) from WT control and LKO female mice on HFD for 10 weeks (n=5). **L**-**O**, Tissue weight (**L**), liver/body weight ratio (**M**), liver TAG content (**N**), liver gross appearance and histology (**O**) from *Clstn3bfl/fl* mice receiving control or *alb-cre* AAV injection on HFD for 10 weeks (n=5). **P**-**Q**, Serum FFA (**P**), cholesterol and TAG level (**Q**) from WT control and LKO mice on HFD for 10 weeks (n=6). **R**-**S**, Serum FFA (**R**), cholesterol and TAG level (**S**) from *Clstn3bfl/fl* mice receiving control or *alb-cre* AAV injection on HFD for 10 weeks (n=6). **T**-**U**, Heat production (**T**) and movement count (**U**) of a WT control and LKO cohort on HFD for 10 weeks over a 45-hour period (n=5). **V**, Daily food intake of WT control and LKO mice on HFD for 10 weeks (n=5). **W**, Body composition analysis of WT control and LKO mice (n=5) on HFD for 10 weeks. **X**, qPCR analysis of hepatic mitochondrial biogenesis and FAO gene expression in WT and LKO mice (n=5) on HFD for 10 weeks. **Y**-**Z**, Fluorescence image (**Y**) and quantitation (**Z**) of FAO blue assay on primary WT and LKO hepatocytes (n=20-24). Scale bar: 100 µm in **F**, **K**, and **O**; 20 µm in the inset of **O**; 5 µm in **Y**. Data are mean ± s.e.m. Statistical significance was calculated by two-way ANOVA with repeated measurement in **T** and **U** and unpaired Student’s two-sided t-test in all other panels.

**Figure S3.**
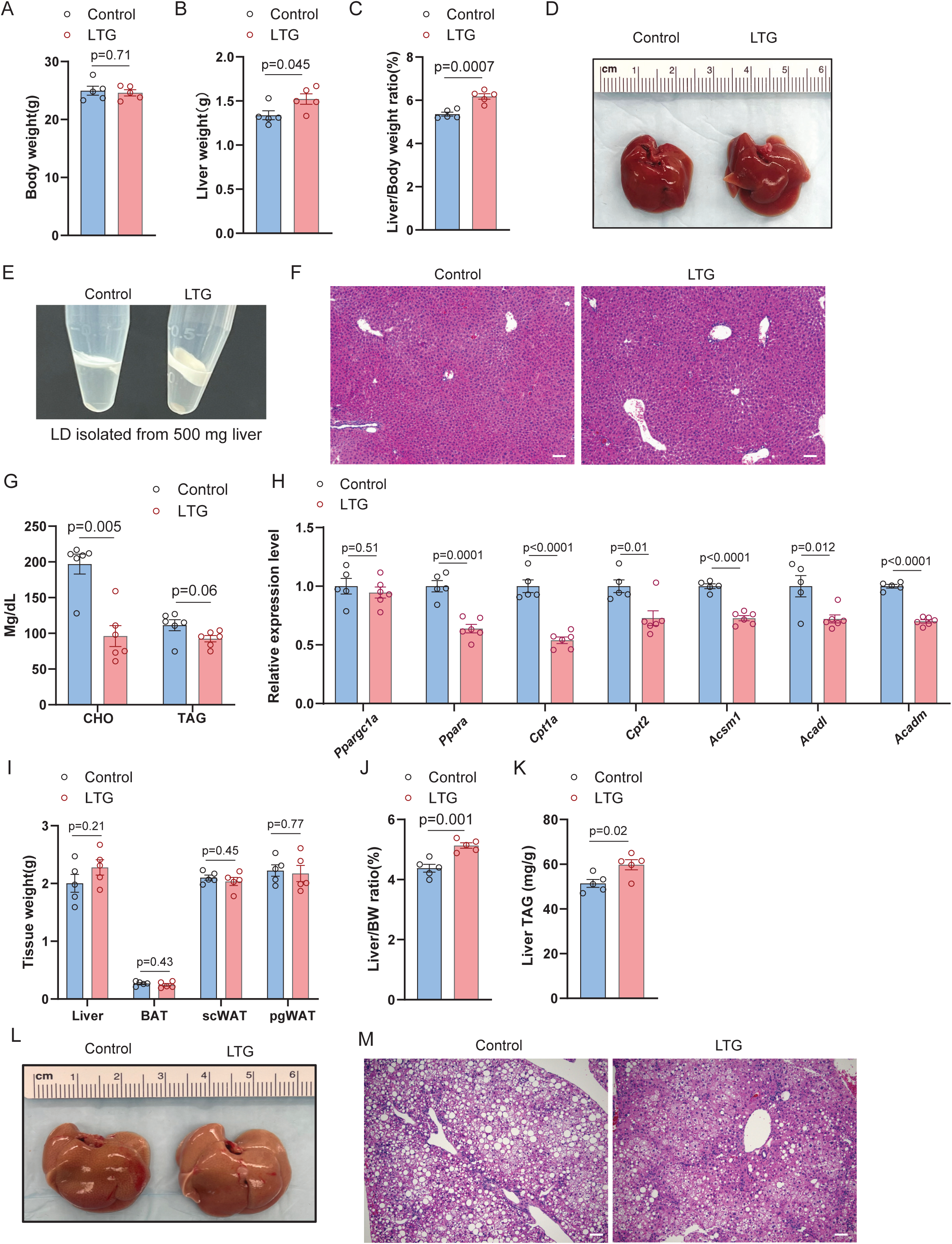
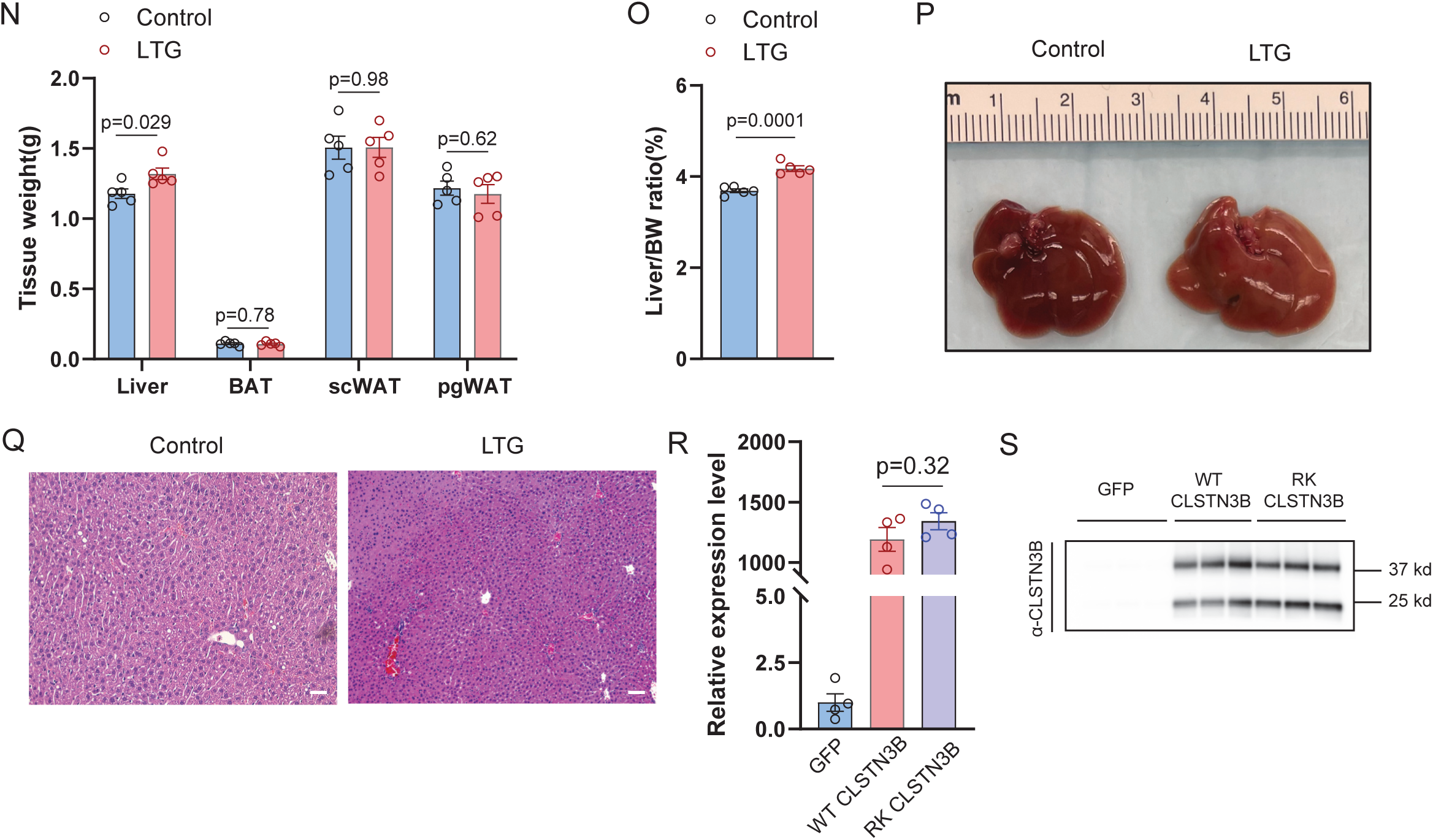
**A**-**F**, Body weight (**A**), liver weight (**B**), liver/body weight ratio (**C**), liver gross appearance (**D**), histology (**E**) and LD layer isolated from 500mg liver tissue (**F**) from male WT control and LTG mice on chow diet (n=5). **G**, Serum cholesterol and TAG level from LTG and control mice on HFD for 10 weeks (n=6). **H**, qPCR analysis of hepatic mitochondrial biogenesis and FAO gene expression from WT control and LTG mice on HFD for 10 weeks (n=5). **I**-**M**, Tissue weights (**I**), liver/body weight ratio (**J**), liver TAG content (**K**), liver gross appearance (**L**), and histology (**M**) from male WT control and LTG mice on WD for 10 weeks (n=5). **N**-**Q**, Tissue weight (**N**), liver/body weight ratio (**O**), liver gross appearance (**P**) and histology (**Q**). **R**-**S**, *Clstn3b* mRNA expression level (**R**) or CLSTN3B protein level (**S**) in liver from mice receiving AAV-GFP, AAV-WT-CLSTN3B or AAV-RK-CLSTN3B injection. Scale bar: 100 µm in **F**, **M**, and **Q**; Data are mean ± s.e.m. Statistical significance was calculated by one-way ANOVA with Tukey’s post hoc test in **R** and unpaired Student’s two-sided t-test in all other panels.

**Figure S4.**
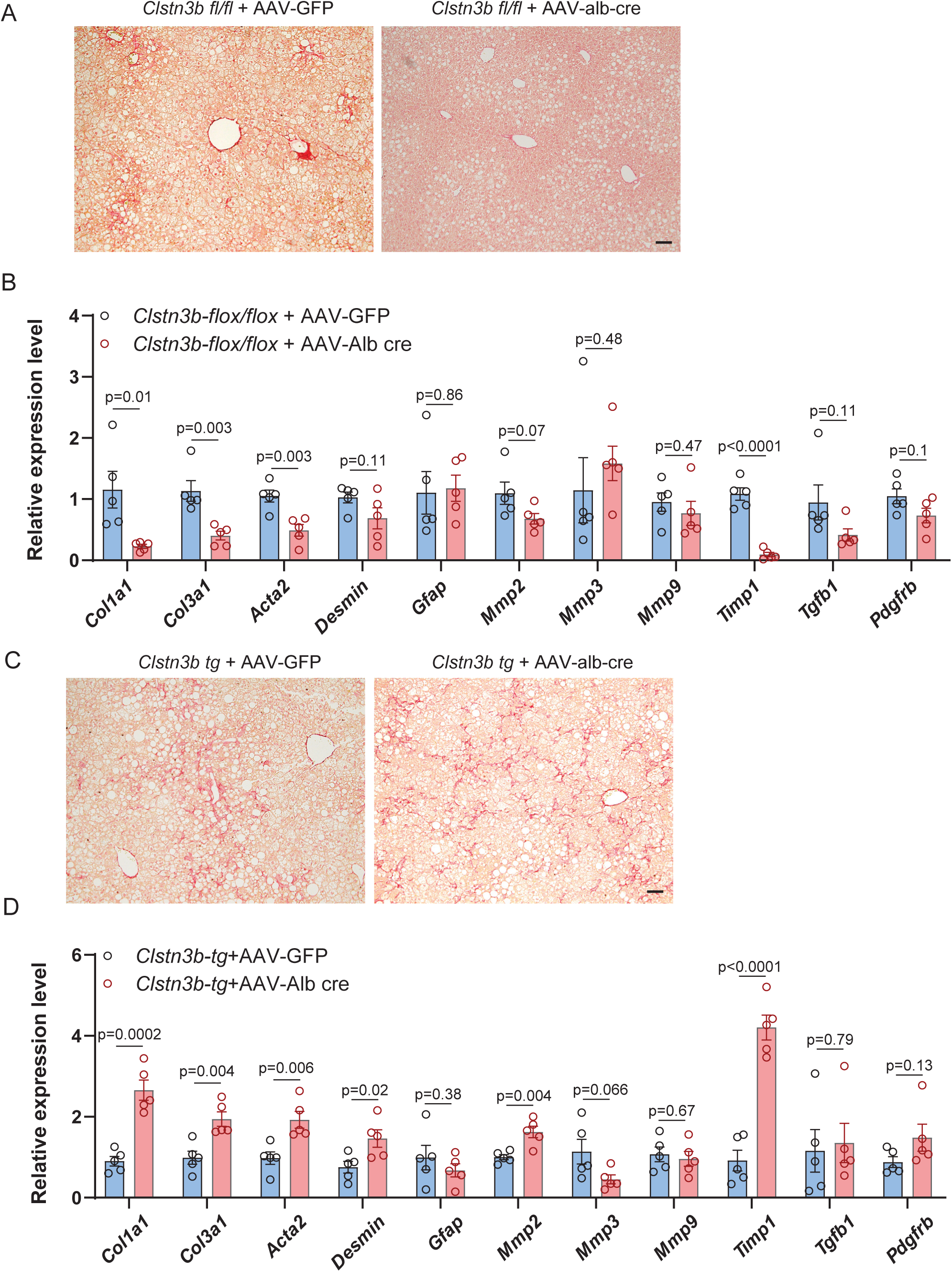
**A**, Sirium Red staining of liver sections of control and AAV-induced liver-specific CLSTN3B KO cohort (n=5) maintained on a WD. **B**, qPCR analysis of genes associated with hepatic fibrosis in the liver of mice (n=5) from **A**. **C**, Sirium Red staining of liver sections of control and AAV-induced liver-specific CLSTN3B transgenic cohort (n=5) maintained on a WD. **D**, qPCR analysis of genes associated with hepatic fibrosis in the liver of mice (n=5) from **c**. Scale bar: 100 µm in **A** and **C**. Data are mean ± s.e.m. Statistical significance was calculated by unpaired Student’s two-sided t-test in all panels.

**Figure S5.**
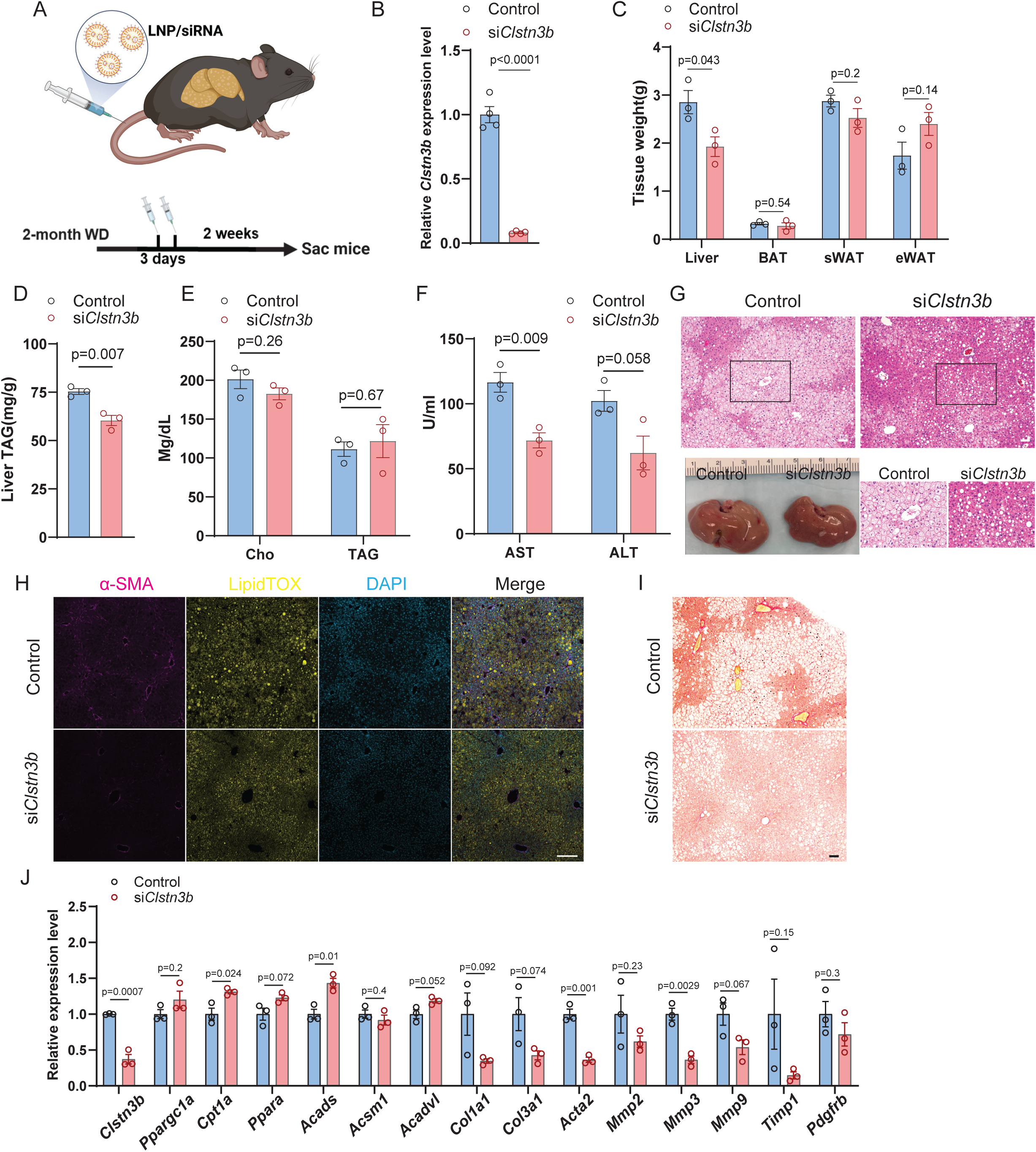
**A**, Schematics of hepatic *Clstn3b* knockdown by LNP/siRNA injection. **B**-**J**, Hepatic *Clstn3b* expression levels (**B**), tissue weights (**C**), liver triglyceride contents (**D**), serum cholesterol and triglyceride levels (**E**), serum AST and ALT levels (**F**), liver gross appearance and histology (**G**), immunostaining of the fibrosis marker α-SMA in liver sections (**H**), Sirium Red staining of liver sections (**I**), and qPCR analysis of hepatic mitochondrial and fibrosis-related gene expression (**J**) from mice receiving control or *Clstn3b*-targeting siRNA injections. Scale bar=100 µm. Data are mean ± s.e.m. Statistical significance was calculated by unpaired Student’s two-sided t-test.

**Figure S6.**
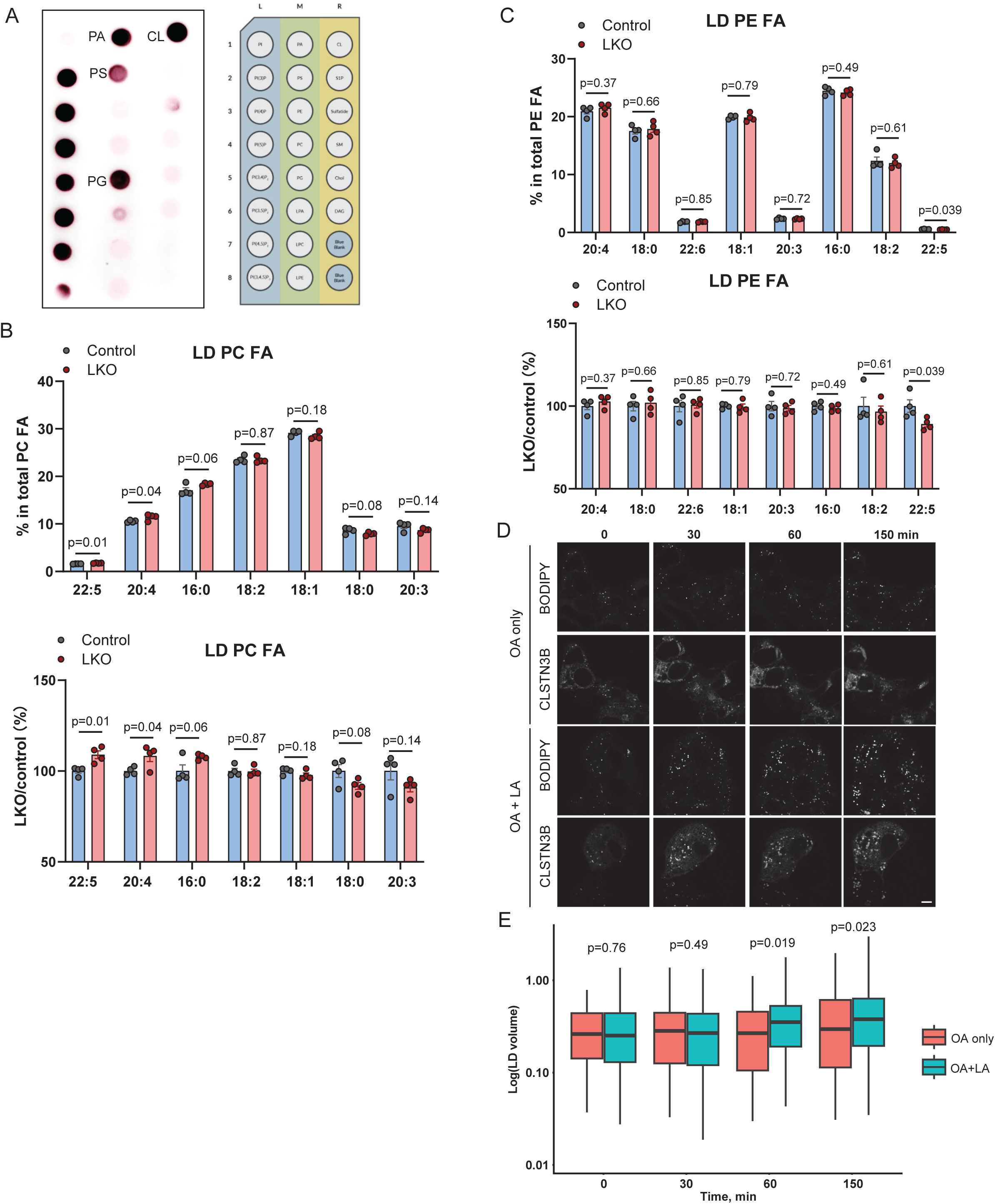
**A**, Lipid strip binding assay with purified recombinant CLSTN3B. Note that CLSTN3B strongly prefers anionic phospholipids. **B**-**C**, PC (**B**) and PE (**C**) acyl chain composition analysis of LD fractions isolated from control of LKO mice after 10-week HFD feeding (n=4). The fraction of each acyl chain type in PC or PE and the relative difference between LKO and control are shown separately. **D**-**E**, Representative images (**D**) and quantitation of LD volume changes over time (**E**) in cells treated with OA alone or OA+LA (n=100-300 for each time point). Data are mean ± s.e.m. Statistical significance was calculated by unpaired Student’s two-sided t-test (**B** and **C**) or Wilcoxon rank-sum test with Benjamini-Hochberg correction (**E**).

**Figure S7.**
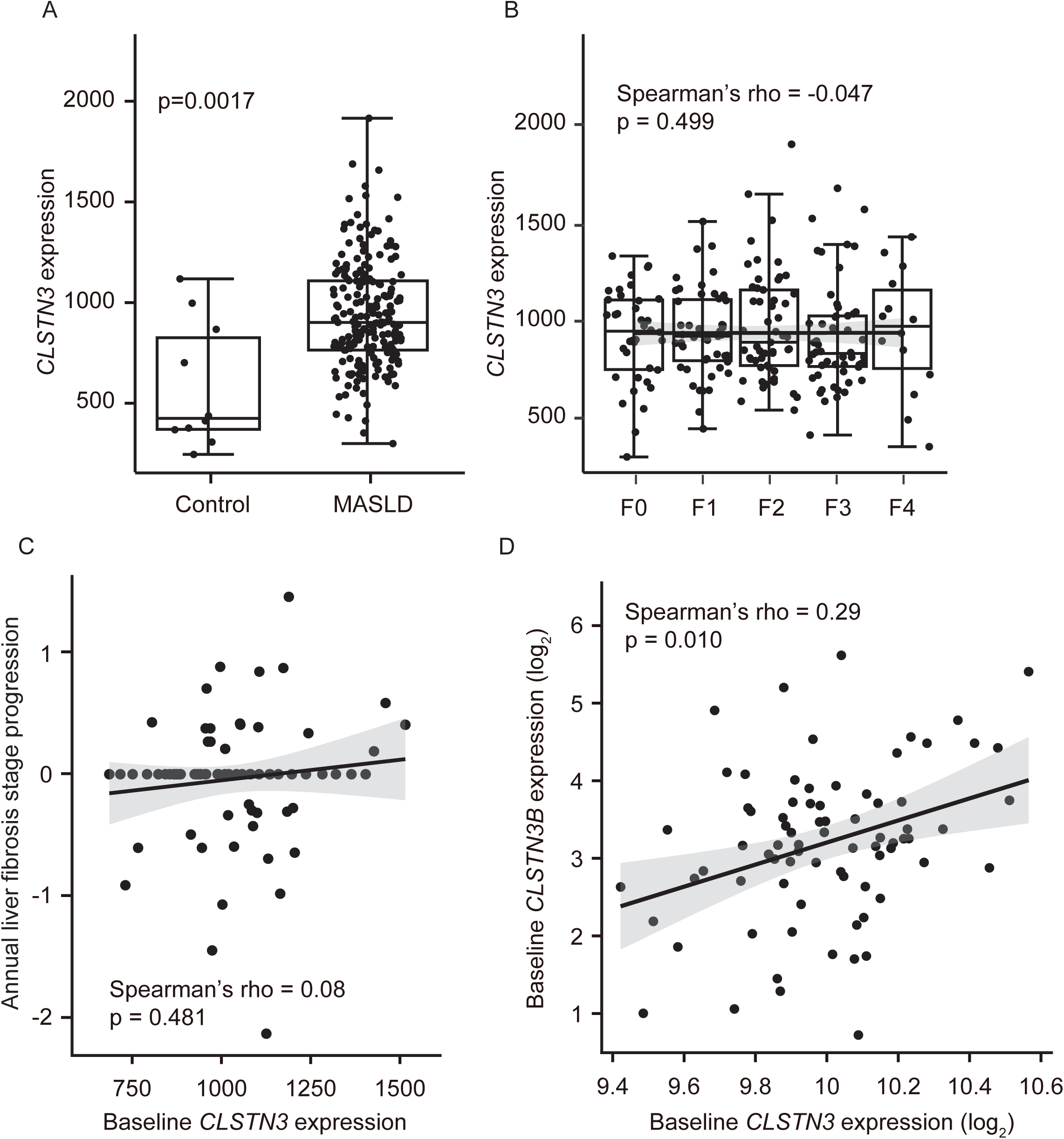
**A**, Hepatic *CLSTN3* expression levels in 206 European MASLD patients compared to 10 healthy obese controls. **B**, Hepatic *CLSTN3* expression levels by liver fibrosis (F) stage in 206 European MASLD patients. **C**, Correlation between baseline *CLSTN3* expression levels and time-adjusted (i.e., annual) progression of liver fibrosis stage in 77 Japanese MASLD patients. **D**, Correlation between baseline *CLSTN3* and *CLSTN3B* expression levels in 77 Japanese MASLD patients. Inter-group difference was assessed by Wilcoxon rank-sum test. Correlation was assessed by Spearman correlation test.

## Notes

### Competing Interest Statement

The authors have declared no competing interest.

